# Multi-modal brain properties are associated with interindividual differences in fear acquisition and extinction

**DOI:** 10.1101/2025.04.05.647350

**Authors:** Christoph Fraenz, Dorothea Metzen, Julian Packheiser, Christian J. Merz, Helene Selpien, Nikolai Axmacher, Erhan Genç

**Affiliations:** Department of Psychology and Neurosciences, Leibniz Research Centre for Working Environment and Human Factors (IfADo), 44139 Dortmund, Germany; Department of Clinical and Biopsychology, Institute of Psychology, Faculty of Educational Sciences and Psychology, TU Dortmund University, 44227 Dortmund, Germany; Department of Social Neuroscience, Institute of Cognitive Neuroscience, Faculty of Psychology, Ruhr University Bochum, 44801 Bochum, Germany; Department of Cognitive Psychology, Institute of Cognitive Neuroscience, Faculty of Psychology, Ruhr University Bochum, 44801 Bochum, Germany; Department of Anesthesiology and Operative Intensive Care Medicine, University Hospital Schleswig-Holstein, 24105 Kiel, Germany; Department of Neuropsychology, Institute of Cognitive Neuroscience, Faculty of Psychology, Ruhr University Bochum, 44801 Bochum, Germany

**Keywords:** Fear acquisition, fear extinction, brain macrostructure, brain microstructure, brain connectivity, functional magnetic resonance imaging

## Abstract

Interindividual differences in fear acquisition and extinction have been related to variation in specific brain correlates. However, variability in experimental setups complicates the integration of findings. Here, we present a combined fear acquisition (n = 101) and extinction (n = 88) experiment in which both phenomena were related to brain correlates obtained via functional magnetic resonance imaging in healthy, young participants. Correlates included regional brain volume, cortical surface area and thickness, neurite density and orientation dispersion, structural and functional connectivity. Fear responses were quantified as changes in skin conductance. Data from 376 brain areas and 70,500 network connections were used as independent variables in regularized regression models. Regression models of fear acquisition could be obtained for all modalities but regional brain volume. There were 284 predictors of which 77 appeared in exactly two models and 19 in exactly three. The latter primarily included brain areas from the somatosensory, insular, cingulate, and frontal cortices. Fear extinction yielded regression models based on neurite density, structural connectivity, and functional connectivity with 112 predictors in total. Two predictors, located in the dorsolateral prefrontal cortex, replicated across exactly two regression models (neurite density and structural connectivity). This study is the first to investigate the neural correlates of both fear acquisition and extinction in an explorative, multi-modal fMRI approach. Results show that numerous brain regions contribute to fear conditioning, some of them via more than one correlate. These findings call for further research to examine the potential interplay between brain correlates shaping fear conditioning.

## 1 Introduction

Fear is a fundamental emotion that plays a crucial role in human survival and adaptation. The acquisition and extinction of fear are essential processes through which individuals learn to respond to and regulate threatening stimuli in their environment (Craske et al., 2006). Fear acquisition refers to the initial learning of fear associations, wherein a neutral stimulus becomes associated with an aversive event, leading to the subsequent elicitation of fear responses. On the other hand, fear extinction involves the gradual reduction and eventual suppression of fear responses when the previously learned fear association is no longer relevant or threatening (Lonsdorf et al., 2017). Individuals show significant differences in their susceptibility to acquiring and extinguishing fear responses (Lonsdorf & Merz, 2017). In recent years, elucidating the neurobiological underpinnings that account for these inter-individual differences in fear acquisition and extinction has been a challenging endeavor for the field of neuroscience. Previous research was able to identify various brain regions in which specific neuronal parameters were associated with an individual’s responsiveness to fear acquisition or extinction. However, potential brain correlates have only been analyzed in isolation of each other. Hence, it remains an open question whether the neurobiology of fear acquisition and extinction is characterized by a multi-modal set of brain properties.

Investigating the neural correlates of fear acquisition and extinction relies on controlled experiments that typically employ classical conditioning paradigms (Lonsdorf et al., 2017). In this process, a neutral stimulus (such as a tone or a visual cue) is paired with an unconditioned stimulus (US, such as a mild electric shock or an aversive image), causing the neutral stimulus to become a conditioned stimulus (CS) that elicits a conditioned fear response. Fear responses can be quantified via physiological measures (such as skin conductance responses, SCRs) and self-report. Fear extinction studies aim to diminish previously established conditioned fear responses by presenting the CS without the aversive event, leading to a gradual reduction in fear.

Numerous studies have explored the neural correlates of fear acquisition and extinction using functional magnetic resonance imaging (fMRI). These investigations have provided valuable insights into the underlying brain regions associated with these processes. For example, the amygdala, a key region involved in emotional processing, has been shown to exhibit heightened activation during fear acquisition, primarily during the early phase of conditioning (Buchel et al., 1999; LaBar et al., 1998; Phelps, 2004). The amygdala’s engagement in fear acquisition has been debated in more recent studies (Visser et al., 2021; Wen et al., 2024; Wen et al., 2022) but is supported by its connectivity with other regions such as the prefrontal cortex (PFC), insula, and hippocampus, which play critical roles in the evaluation, regulation, and contextualization of fear responses (Milad & Quirk, 2012; but see Fullana et al., 2019). Milad, Wright, et al. (2007) utilized fMRI to demonstrated that the ventromedial PFC is deactivated during fear acquisition but activated during fear extinction. In addition, results showed that ventromedial PFC activation during extinction recall is positively correlated to extinction retention. Recent meta-analyses have identified whole brain networks showing increased activity in response to fear-or safety-related stimuli (Fullana et al., 2018; Fullana et al., 2016).

While much research has focused on task-based brain activity that is exhibited during fear acquisition and extinction, recent studies have started exploring the relationship between resting-state brain connectivity and fear conditioning. For example, it has been demonstrated that the process of fear conditioning can alter the resting-state connectivity between various brain regions. (Feng et al., 2014; Feng et al., 2013; Hermans et al., 2017; Martynova et al., 2020; Schultz et al., 2012). Respective connections were, among others, constituted by the amygdala, dorsal anterior cingulate cortex, ventromedial PFC, insula, and hippocampus. These regions have been related to fear acquisition and/or extinction in previous task-based studies. Resting-state brain connectivity is likely to change as the result of fear conditioning but it can also, at least to some extent, predict the effect of fear conditioning. For example, it has been shown that resting-state connectivity between amygdala and ventromedial PFC, measured after fear acquisition, is positively correlated with the effect of subsequent fear extinction (Feng et al., 2016). These findings suggest that resting-state brain connectivity holds promise as a potential biomarker for individual differences in fear acquisition, extinction, and susceptibility to fear-related disorders.

In addition to task-based and resting-state fMRI studies, previous studies have explored the structural correlates of fear acquisition and extinction using structural MRI techniques. For instance, cortical thickness of the posterior insula (Hartley et al., 2011) and the dorsal anterior cingulate cortex (Milad, Quirk, et al., 2007) has been shown to be positively associated with fear responses evoked in a classical fear conditioning paradigm. Following a similar approach, a recent study investigating fear learning in anxiety patients and healthy controls found cortical thickness of the dorsomedial and dorsolateral PFC to be negatively associated with SCRs (Abend et al., 2020). In view of subcortical regions, it has been demonstrated that left amygdala volume is positively correlated with the magnitude of conditioned fear responses, whereas bilateral hippocampal volumes are associated with the magnitude of differential contingency ratings, i.e., how well participants can remember which CS was paired with an US and which was not (Cacciaglia et al., 2015). Furthermore, it has been shown that larger hippocampal volume is indicative of stronger SCRs to CS+ presentations during fear acquisition (Pohlack et al., 2012). Right amygdala volume was found to be positively correlated with SCR diffences between CS+ and CS-presentations during fear acquisition (Winkelmann et al., 2016).

In addition to studying brain macrostructure and functional brain connectivity, previous research has employed diffusion tensor imaging (DTI) and fiber tracking techniques to investigate the role of structural brain connectivity in fear learning processes. DTI allows the examination of white matter tracts in the brain, providing insights into the integrity and organization of neural connections. For instance, fractional anisotropy (FA), a marker of white matter integrity, has been associated with fear extinction (Nees et al., 2019) and fear renewal (Hermann et al., 2017) expression. Higher FA values measured in the hippocampal cingulum were indicative of higher SCRs during the extinction of contextual conditioned fear responses as well as stronger fear renewal. Furthermore, it has been demonstrated that patients suffering from post-traumatic stress disorder and trauma-exposed controls show significant differences with regard to white matter integrity of specific brain regions (Kunimatsu et al., 2020). In another study comparing trauma-exposed individuals and trauma-unexposed individuals, DTI was used to extract local, connectivity, and network features of fear-circuitry related brain regions, including the amygdala, orbitofrontal and ventromedial PFC, hippocampus, insula, and thalamus. By analyzing the DTI derived measures via a machine learning approach, the authors were successful in distinguishing between the two groups at each stage of recovery (Im et al., 2017).

As our understanding of the neural correlates of fear acquisition and extinction continues to evolve, researchers have been exploring advanced MRI techniques to investigate the microstructural properties of the brain. Alongside the previously discussed methods such as task-based and resting-state fMRI, regional brain volume, cortical thickness, and structural connectivity measures derived from DTI, there is another promising diffusion MRI technique called Neurite Orientation Dispersion and Density Imaging (NODDI) (Zhang et al., 2012). NODDI provides valuable insights into the microstructural correlates of fear acquisition and extinction by characterizing the density of neurites (axons and dendrites) and the dispersion of their orientations. While NODDI is a relatively novel technique that has not been applied to investigate human fear acquisition and extinction, its potential for quantifying microstructural alterations in the brain has been utilized in other fields, e.g., to investigate interindividual differences in cognitive ability (Genc et al., 2018), language processing (Ocklenburg et al., 2018), and hemispheric asymmetries (Schmitz et al., 2019).

Integrating findings from multiple neuroimaging techniques, including functional and structural measures as well as connectivity analyses, allows for a comprehensive examination of the neural correlates of fear acquisition and extinction. Such multi-modal approaches hold promise in unraveling the complex interactions and mechanisms underlying fear-related behaviors. However, it is challenging to integrate results from previous studies given their diverse approaches. Typically, these studies are focused on only one isolated neuroimaging technique (either structural, functional, or diffusion MRI). In most cases, they investigate merely one specific phase of fear conditioning (either fear acquisition or extinction). Moreover, many studies only analyze specific brain regions, usually those that have been related to fear learning in previous research. Consequently, they run the risk of missing any significant effects outside of this a priori defined set. Finally, different studies may use different methods (SCRs, pupil dilation, heart rate, self-reports) and statistics (reaction towards CS+ or CS-alone, difference between CS+ and CS-reactions) to quantify fear responses.

Here, we investigated the underpinnings of both fear acquisition and extinction, allowing for a comprehensive examination of these distinct processes within a single research effort. We combined functional and structural connectivity analyses as well as macrostructural markers such as cortical thickness, cortical surface area, and regional brain volume. We also included NODDI as a novel technique to quantify the microstructural correlates of fear acquisition and extinction. The recruited sample was comparatively large, including approximately 100 individuals. Our analyses followed a whole-brain approach, utilizing a well-established parcellation scheme from the Human Connectome Project. By encompassing multiple modalities, investigating both fear acquisition and extinction, and employing a whole-brain analysis, our study is capable of exploring a variety of neural mechanisms potentially underlying fear learning in humans.

## 2 Methods

### 2.1 Participants

The sample recruited for the current study consisted of 138 young and healthy participants, most of them college students. Several participants had to be excluded from the initially recruited sample due to various reasons: Two participants were removed because of incomplete or corrupted MRI data files. Thirteen participants reported a CS-/US contingency that was equal to or larger than their reported CS+/US contingency, indicating absent contingency awareness, which is a necessary prerequisite to observe conditioned SCRs (Tabbert et al., 2011). Finally, 22 participants had to be removed from the analysis of fear acquisition data and 35 participants had to be removed from the analysis of fear extinction data because of technical difficulties with the electrical stimulation, corrupted or missing SCR data files, or excessive noise in the participants’ SCR recordings. Hence, all analyses related to fear acquisition were carried out with data from 101 participants (56 women; 18 to 26 years, M = 21.97, SD = 2.07), while analyses related to fear extinction included data from 88 participants (47 women; 18 to 26 years, M = 22.08, SD = 2.11). Even though these sample sizes are comparatively large for a neuroimaging study concerned with fear learning, it has to be noted that the number of participants (observations) was vastly exceeded by the number of predictors in our regression analyses. However, we addressed this issue by employing elastic-net regularization which has been found to be the superior form of variable selection over other methods such as lasso regularization. For example, Zou & Hastie (2005) carried out regression analyses with elastic-net regularization in a data set with 7129 predictors and 72 observations.

We did not observe significant age differences between male and female participants, neither in the fear acquisition group (t(99) = 1.696, p = .0931) nor in the fear extinction group (t(86) = 1.505, p = .136). In order to control for confounding effects caused by handedness, only right-handed individuals, as measured by the Edinburgh Handedness Inventory (Oldfield, 1971), were recruited. All participants had normal or corrected-to-normal vision and were able to understand the provided written and oral instructions. They were either paid for their participation or received course credit. All participants were naive to the purpose of the study and had no former experience with the fear acquisition and extinction paradigms used for the experiment. Participants reported no history of psychiatric or neurological disorders and matched the standard inclusion criteria for fMRI examinations. The study was approved by the local ethics committee of the Faculty of Psychology at Ruhr University Bochum (8^th^ of November 2016, application number 327). All participants provided written informed consent prior to participation and were treated in accordance with the Declaration of Helsinki.

### 2.2 Acquisition of Imaging Data

All imaging data were acquired at Bergmannsheil hospital in Bochum (Germany) using a 3 Tesla Philips Achieva scanner with a 32-channel head coil. Scanning included T1-weighted imaging, diffusion-weighted imaging, and resting-state imaging. The T1-weighted and diffusion-weighted images were obtained on day 1, whereas the resting-state images were obtained directly before the fear acquisition and fear extinction paradigm on day 2 (Figure 1).

**Figure 1.**
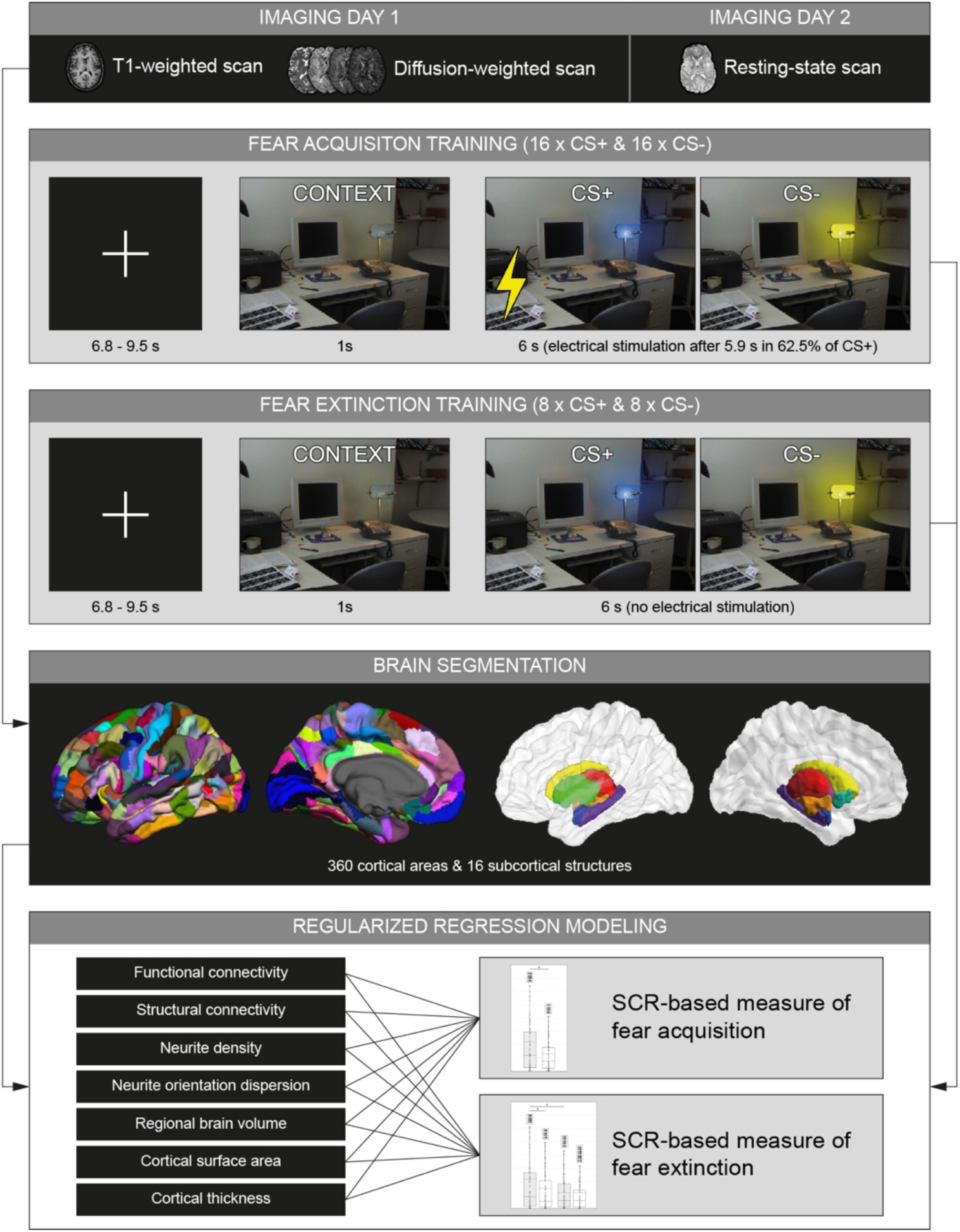
Schematic depiction of data acquisition, preprocessing, and analysis. On the first day, participants underwent T1-weighted and diffusion-weighted imaging. On the second day, they participated in an fMRI resting-state scan that was followed by a fear acquisition and extinction paradigm. At the beginning of each trial, a white fixation cross was presented on a black background for 6.8 - 9.5 seconds. Next, the context image was presented for 1 second, which was followed by the presentation of the CS+ (blue lamplight) or the CS-(yellow lamplight) for another 6 seconds. During the fear acquisition paradigm, the CS+ was paired with electrical stimulation as the US (indicated by a yellow bolt) in 62.5 % of trials administered 5.9 seconds after CS+ onset. CS+ and CS-were presented 16 times each in pseudo-randomized order. During the fear extinction paradigm, CS+ and CS-were presented 8 times each but without US. All brain images obtained prior to the fear acquisition and extinction paradigm were parcellated into 360 cortical and 16 subcortical regions. T1-weighted images were used to compute the regional brain volume, cortical surface area, and cortical thickness of each region. Diffusion-weighted images were used to compute measures of neurite density and orientation dispersion for each region. Moreover, fiber tracking was performed to estimate structural connectivity between all possible combinations of brain regions. Likewise, fMRI resting-state scans were used to compute average BOLD signal time courses for each region. These were correlated with each other to obtain estimates of functional connectivity between all possible combinations of brain regions. Fear acquisition was quantified by subtracting the average SCR to CS-trials from the average SCR to CS+ trials. In case of fear extinction, CS+ and CS-responses were averaged across the first four and the last four trials, respectively. The difference between average CS+ and CS-responses from the second half was subtracted from the difference between average CS+ and CS-responses from the first half. For each type of brain imaging metric, two regularized regression models were computed. One with the SCR-based measure of fear acquisition as its dependent variable and one with the SCR-based measure of fear extinction.

#### 2.2.1 T1-weighted Imaging

For the purpose of brain image parcellation, and in order to quantify macrostructural parameters such as regional brain volume, cortical thickness, and cortical surface area, T1-weighted high-resolution anatomical images were acquired (MP-RAGE, TR = 8.2 ms, TE = 3.7 ms, flip angle = 8°, 220 slices, matrix size = 240 mm x 240 mm, resolution = 1 mm x 1 mm x 1 mm). Scanning time was around 6 minutes.

#### 2.2.2 Diffusion-weighted Imaging

For the purpose of fiber tracking and the analysis of NODDI coefficients, diffusion-weighted images were acquired using echo planar imaging (TR = 7652 ms, TE = 87 ms, flip angle = 90°, 60 slices, matrix size = 112 x 112, voxel size = 2 mm x 2 mm x 2 mm). Diffusion weighting was based on a multi-shell, high angular resolution scheme consisting of diffusion-weighted images for b-values of 1000, 1800, and 2500 s/mm^2^, respectively, applied along 20, 40, and 60 uniformly distributed directions. All diffusion directions within and between shells were generated orthogonal to each other using the MASSIVE toolbox (Froeling et al., 2016). Additionally, eight volumes with no diffusion weighting (b = 0 s/mm²) were acquired as an anatomical reference for motion correction and computation of NODDI coefficients. Diffusion-weighted data was collected with reversed phase-encode directions, resulting in pairs of images with distortions going in opposite directions. Scanning time was around 18 minutes.

#### 2.2.3 Resting-state Imaging

For the analysis of functional brain connectivity, fMRI resting-state images were acquired prior to fear acquisition using echo planar imaging (TR = 2500 ms, TE = 30 ms, flip angle = 90°, 40 slices, matrix size = 112 mm x 112 mm, resolution = 2 mm x 2 mm x 3 mm). Participants were instructed to close their eyes, relax, and think of nothing in particular. Scanning time was around 8 minutes.

### 2.3 Fear Acquisition and Extinction Paradigm

On day 2, all participants completed a differential fear acquisition paradigm followed by a fear extinction paradigm while lying in the fMRI scanner. The stimuli and procedures for these paradigms were modified from Milad, Wright, et al. (2007). Participants were instructed to pay close attention to the images presented during fear acquisition and extinction. They were also told that electrical stimulation may or may not be presented during the experiment. Electrical stimulation (1 ms pulses with 50 Hz for a duration of 100 ms) was applied as the US via a constant voltage stimulator (STM2000 BIOPAC systems, CA, USA) along with two electrodes attached to the fingertips of the first and second fingers of the right hand. The intensity of electrical stimulation was adjusted for each participant individually prior to the resting-state scan preceding the fear acquisition and extinction paradigm on day 2. For this purpose, electrical stimulation was administered at 30 V and raised in increments of 5 V until participants rated the sensation as very unpleasant but not painful. The stimuli used for fear acquisition and extinction were presented using the Presentation software package (Neurobehavioral Systems, Albany, CA) and MR compatible LCD-goggles (Visuastim Digital, Resonance Technology Inc, Northridge, CA). Both paradigms featured a picture of an office room with a switched off desk lamp as the context image. CS presentation was implemented by the desk lamp lighting up in either blue (CS+) or yellow (CS-). At the beginning of each trial, a white fixation cross was presented on a black background for 6.8 - 9.5 seconds. Next, the context image was presented for 1 second, which was immediately followed by the presentation of CS+ or CS-for another 6 seconds. During fear acquisition, the CS+ was paired with electrical stimulation in 62.5 % of trials (10 out of 16). The stimulation was administered 5.9 seconds after CS+ onset and co-terminated with CS+ offset. Exemplary trials for fear acquisition and extinction are depicted in Figure 1. Fear acquisition included a total of 32 trials with CS+ and CS-being presented 16 times each in pseudo-randomized order, while fear extinction included 16 trials with 8 CS+ and 8 CS-presentations, respectively. The first two trials always consisted of one CS+ and one CS-presentation and so did the last two trials. During fear acquisition, the first and last CS+ presentations were always paired with electrical stimulation. There were no trials that presented the same type of CS more than twice in consecutive order. CS+ and CS-presentations were distributed equally across both halves. SCRs were assessed using two Ag/AgCl electrodes filled with isotonic (0.05 NaCl) electrolyte medium placed on the hypothenar eminence right below the fifth finger of the left hand. Data were recorded with a sampling rate of 5,000 Hz using Brain Vision Recorder software (Brain Products GmbH, Munich, Germany).

Subsequent to fear acquisition and extinction, participants had to rate the contingencies between CS+, CS-, and US. First, they were asked to estimate the number of electrical stimulations they had received during the respective paradigm. If they reported to have received at least one electrical stimulation, which was to be expected after fear acquisition, participants were also asked to rate the unpleasantness of the last electrical stimulation on a 9-point Likert scale (1 - “not unpleasant”, 9 - “very unpleasant”) and to estimate at what rates the blue and yellow lamplights had been followed by an electrical stimulation. If participants reported to have received no electrical stimulation at all, which was to be expected after fear extinction, they were merely asked if they had noticed any differences between the two images and how often the blue and the yellow lamplights were presented.

### 2.4 Preprocessing of Imaging Data

#### 2.4.1 Preprocessing of T1-weighted Data

For the purpose of reconstructing the cortical surfaces of T1-weighted images, we used published surface-based methods in FreeSurfer (http://surfer.nmr.mgh.harvard.edu, version 6.0.0) along with the CBRAIN platform (Sherif et al., 2014). The details of this procedure have been described elsewhere (Dale et al., 1999; Fischl et al., 1999). The automatic reconstruction steps included skull stripping, gray and white matter segmentation as well as reconstruction and inflation of the cortical surface. These preprocessing steps were performed for each participant individually. Subsequently, each individual segmentation was quality-controlled slice by slice and inaccuracies of the automatic steps were corrected by manual editing if necessary.

The analyses of structural and functional brain connectivity, macrostructural brain properties, and NODDI coefficients followed a region-based approach, for which we considered brain regions as defined by the Human Connectome Project’s multi-modal parcellation (HCPMMP) (Glasser et al., 2016) and FreeSurfer’s automatic subcortical segmentation (Figure 1). The HCPMMP delineates 180 cortical brain regions per hemisphere and is based on the cortical architecture, function, connectivity, and topography of 210 healthy individuals. From FreeSurfer’s automatic subcortical segmentation, we included a set of eight subcortical regions per hemisphere (thalamus, caudate nucleus, putamen, pallidum, hippocampus, amygdala, accumbens area), various ventricle masks (lateral ventricle, inferior lateral ventricle, third ventricle, fourth ventricle), and a mask covering the whole cerebral white matter compartment (Fischl et al., 2002). In total, 360 cortical masks, 16 subcortical masks, 8 ventricle masks, and one white matter mask were selected and linearly transformed into the native spaces of the diffusion-weighted and resting-state images to guide further analyses.

#### 2.4.2 Preprocessing of Diffusion-weighted Data

Diffusion-weighted data were prepared for fiber tracking and the computation of NODDI coefficients via a preprocessing pipeline comprising the following steps. First, images were corrected for signal drift (Vos et al., 2017) using ExploreDTI (Leemans et al., 2009). Second, we utilized the topup command from the FSL toolbox (Smith et al., 2004) to estimate the susceptibility-induced off-resonance field based on pairs of images with opposite phase-encode directions (see section 2.3.2 Diffusion-weighted Imaging). Third, the topup output was used in combination with the eddy command (Andersson & Sotiropoulos, 2016), which is also part of the FSL toolbox, to correct for susceptibility, eddy currents, and head movement. Importantly, we also performed outlier detection during this step to identify slices where signal had been lost due to head movement during the diffusion encoding (Andersson et al., 2016).

As described above, 376 cortical and subcortical regions were transformed into the native space of the diffusion-weighted images (see section 2.4.1 Preprocessing of T1-weighted Data). We utilized these regions as seed and target masks for probabilistic fiber tractography. We employed a dual-fiber model as implemented in the latest version of BEDPOSTX. The model allows for the representation of two fiber orientations per voxel when more than one orientation is supported by the data. This enables modelling of crossing fibers and produces more reliable results compared with single-fiber models (Behrens et al., 2007). Probabilistic fiber tractography was carried out using the classification targets approach implemented in FMRIB’s Diffusion Toolbox (Behrens et al., 2003; Cohen et al., 2009). At each voxel, 5,000 tract-following samples were generated with a step length of 0.5 mm and a curvature threshold of 0.2 (only allowing for angles larger than 80°). The connectivity between a seed voxel and a specific target region was quantified as the number of streamlines originating in the seed voxel and reaching the respective target region. The connectivity between two brain regions was computed as the sum of all streamlines proceeding from the seed to the target region and vice versa. By doing so, we obtained a structural connectome for each participant, comprising 376 network nodes connected via a total of 70,500 unique network edges.

For the purpose of extracting NODDI coefficients from the preprocessed diffusion-weighted data, we utilized the AMICO toolbox (Daducci, Canales-Rodriguez, et al., 2015). The AMICO approach is based on a convex optimization procedure that converts the non-linear fitting, which is usually applied as part of the NODDI technique, into a linear optimization problem (Daducci, Dal Palu, et al., 2015). This reduces processing time dramatically (Sepehrband et al., 2016). The NODDI technique features a three-compartment model distinguishing intra-neurite, extra-neurite, and CSF environments. For this study, we decided to focus on the intra- and extra-neurite compartments exclusively. Intra-neurite environments are characterized by a stick-like or cylindrically symmetric diffusion signal that is created when water molecules are restricted by the membranes of neurites. In white matter, this kind of diffusion is likely to emerge from the presence of axons. In gray matter, it serves as an indicator of dendrites and axons forming the neuropil. For each voxel, NODDI estimates the intra-neurite volume fraction (INVF), i.e., the portion of the diffusion signal that can be attributed to intra-neurite environments (Jespersen et al., 2010; Jespersen et al., 2012; Zhang et al., 2012). This measure is used to quantify neurite density. In addition, NODDI also provides a tortuosity measure coupling the intra- and the extra-neurite spaces. This metric, known as the neurite orientation dispersion index (ODI), reflects the alignment or dispersion of axons in white matter or axons and dendrites in gray matter (Billiet et al., 2015; Zhang et al., 2012). For each participant, we computed mean values of INVF and ODI for all cortical and subcortical regions (see section 2.4.1 Preprocessing of T1-weighted Data).

#### 2.4.3 Preprocessing of Resting-state Data

Resting-state data were preprocessed using MELODIC, which is part of the FSL toolbox. The preprocessing involved several steps: Discarding the first two volumes from each resting-state scan to allow for signal equilibration, motion and slice-timing correction, and high-pass temporal frequency filtering (0.005 Hz). Spatial smoothing was not applied in order to avoid the introduction of spurious correlations in neighboring voxels.

For each cortical and subcortical region (see section 2.4.1 Preprocessing of T1-weighted Data), 376 in total, we calculated a mean resting-state time course by averaging the preprocessed time courses of corresponding voxels. Next, we computed partial correlations between the average time courses of all pairs of brain regions, while controlling for several nuisance variables. We regressed out the trajectories of 6 head motion parameters as well as the mean time courses extracted from the white matter and ventricle masks (Fraenz et al., 2021). The resulting partial correlation coefficients were subjected to a Fisher z-transformation (Fisher, 1921), which produced normally distributed data suitable for further testing. By doing so, we obtained a functional connectome for each participant, comprising 376 network nodes connected via a total of 70,500 unique network edges.

### 2.5 Preprocessing of Skin Conductance Responses

The preprocessing of SCR data was carried out via the Psychophysiological Modelling toolbox in MATLAB (PsPM, version 6.0.0). First, raw data were trimmed and filtered using a median filter with varying amount of time points, such that an optimal number could be found for each participant. Subsequently, artifacts in the trimmed and filtered data were detected using a script based on the automated quality assessment procedure developed by Kleckner et al. (2018). Next, each dataset was manually checked for residual artifacts that may have been missed by the automated artifact detection tool. If any were found, PsPM was used to manually mark all remaining artifacts for subsequent removal. This step was carried out by three researchers with extensive experience in preprocessing SCR data. Participants with an excessive number of artifacts in their data were excluded from further statistical analysis (see section 2.1 Participants).

After preprocessing, artifact-free data were analyzed with Dynamic Causal Modelling (DCM), as this approach is highly indicated when CS presentation is longer than 0.75 seconds (Bach et al., 2010). In order to ensure accurate estimation of conditioned and unconditioned fear responses, only trials with less than a total of 6 seconds of missing epochs (after data cleaning, see above) were included in the analysis. For this reason, some trials could not be estimated due to the presence of an excessive number of missing values. For both fear acquisition and extinction, we modelled the conditions “Context” (fixed event), “CS onset” (fixed event), “CS interval” (flexible event), and “US” (fixed event) (Bach et al., 2010). It is important to note that PsPM assumes the event sequence to be the same for all trials. Hence, in case of non-reinforced trials (no US), we modelled US omissions by adding an event of the same duration as the actual US in the place where the shock would have occurred during reinforced trials. As it is possible that the response to the CS+ is affected by the response to the US (due to overlap in their timings at the end), we applied two additional steps. First, we gradually decreased the time interval between CS+ onset and US onset, to the point that there would be no differences between CS+US+ and CS+US-trials. Second, for each trial we ensured that the onset of the Gaussian bump plus at least two standard deviations did not overlap with the onset of the US. Both procedures ensured that the response to the CS+ during reinforced trials was not contaminated by the response to the US. Indeed, we found the difference between CS+US+ and CS+US-trials not to be significant, suggesting that our method appropriately disambiguated both types of response. SCR data obtained during fear acquisition and extinction were normalized in the same way and analyzed via the same DCM approach to achieve a high degree of comparability. Once modelling was completed, the results of each trial were manually inspected to ensure a proper model fit. Finally, for each participant, we extracted the amplitude for each trial and for each phase of the experiment. These data, referred to as SCR values in the following, were subjected to statistical analysis.

### 2.6 Statistical Analysis

All statistical analyses were either conducted in MATLAB (version R2022b) or R Studio (version 1.3.1093) with R (version 4.1.0). In order to quantify the effect of fear acquisition, we computed the difference between mean SCR values of CS+ and CS-trials (Supplementary Figure 1, left). Here, a large difference should indicate a strong effect of fear acquisition, assuming that participants show a more pronounced SCR towards CS+ but not CS-. Ideally, all 16 CS+ and CS-trials (see section 2.3 Fear Acquisition and Extinction Paradigm) from a participant’s fear acquisition were averaged to generate these mean SCR values. However, in some trials, recordings were too noisy to be used for averaging. If this was the case, all remaining CS+ and CS-trials were used to generate a participant’s mean SCR value. We analyzed the overall effect of fear acquisition by conducting a one-sample t-test comparing the mean CS+ values of all participants against all mean CS-values.

SCR during fear extinction is typically characterized by high amplitudes during early CS+ trials which decrease over time. In contrast, SCR towards CS-remains relatively flat across all trials. Given this pattern, we decided to use a different measure to quantify the effect of fear extinction. In detail, we computed the difference between mean SCR values of CS+ and CS-trials for the first and second half of fear extinction separately. Each of these mean values was obtained by averaging across four trials, given that no trials had to be omitted due to an excessive number of missing epochs (see 2.5 Preprocessing of Skin Conductance Responses). We subtracted the difference computed for the second half from that computed for the first half (Supplementary Figure 1, right). This difference between differences served as an indicator of how strongly a participant’s initial fear response decreased over time when no electrical stimulation was administered. More precisely, a strong effect of fear extinction can be assumed if the difference between mean CS+ and CS-values, which is typically high for the first half of fear extinction, shows a significant decrease and approaches 0 for the second half. We tested this interaction between stimulus type (CS+ or CS-) and time (first or second half of fear extinction) by conducting a two-way ANOVA with repeated measures.

To examine the relationship between neural and measures of fear acquisition or extinction, we utilized the cv.glmnet function from R’s glmnet package to perform regularized regression analysis. Unlike traditional regression methods, this approach does not rely on *p*-values to establish the statistical significance of an independent variable and thus avoids the need for a conventional correction procedure for multiple comparisons such as FDR or Bonferroni (Gongora et al., 2020). Instead, regularized regression analysis applies a penalty to effect sizes through a form of regularization. This approach shrinks small effects towards zero, leaving only strong effects as non-zero. For a detailed explanation of this method, refer to Zou and Hastie (2005) as well as Serang et al. (2017). In general, the model includes all independent variables and subjects their corresponding regression weights to penalization. The lambda penalty term is calculated using k-fold cross validation to prevent overfitting. In this process, the data is divided into k subsets, with one subset designated as the testing-set and the remaining subsets used as the training-set. This is repeated k times, with each subset serving as the testing-set once. Following the penalization procedure, all independent variables with non-zero effects are selected for inclusion. Here, we employed elastic-net regression analysis, a type of regularized regression that combines lasso and ridge regression (Zou & Hastie, 2005). Ridge and lasso regression differ in that lasso regression can shrink a parameter to zero, while ridge regression can only asymptotically shrink a parameter toward zero. Lasso regression is suitable for models with many variables that have no or minimal effect on the dependent variable, while ridge regression is best suited for models in which most variables significantly impact the dependent variable. Elastic-net regression is the preferable approach when no clear expectations exist for every variable. Moreover, it is better than lasso regression at handling correlations between variables (Zou & Hastie, 2005). In previous research, elastic-net regression has been successfully applied to investigate the association between intelligence and the microstructure of white-matter tracts (Gongora et al., 2020) as well as the link between polygenetic scores, brain connectivity, and intelligence (Genç et al., 2023).

For each regression model, the number of cross-validation folds was set to k = 10 and α was set to α = 0.5. A measure of fear acquisition or extinction served as the dependent variable, while the independent variables were constituted by the values from one of the seven brain parameters. In more detail, the regression models for resting-state connectivity (RSC) and diffusion-weighted imaging connectivity (DWIC) both included 70,500 independent variables, one for each connection from the overall network (see section 2.4.2 Preprocessing of Diffusion-weighted Data and section 2.4.3 Preprocessing of Resting-state Data). The regression models for regional brain volume (VOL), INVF, and ODI each included 376 independent variables, namely 360 for all regions of interests (ROIs) from the HCPMMP and 16 for all subcortical ROIs. The regression models for cortical surface area (SURF) and cortical thickness (THICK) both only comprised 360 independent variables since respective parameters were not available for subcortical regions. In all models, sex and age were used as control variables. Finally, we checked whether the seven regression models, featuring either fear acquisition or extinction data, showed any overlap with regard to the predictors selected by elastic-net analysis. To this end, we considered all relevant ROIs exhibited by the SURF, VOL, THICK, INVF, and ODI regression models as well as all brain regions (network nodes) constituting relevant connections in the RSC and DWIC regression models.

## 3 Results

### 3.1. SCR-based outcome of fear acquisition

Mean SCR values from CS+ and CS-presentations during fear acquisition were compared with each other using a one-sample t-test. Results showed that SCR was significantly higher towards CS+ than CS-(t(100) = 3.162, p < .01), indicating that fear acquisition was successful (Figure 2, left).

**Figure 2.**
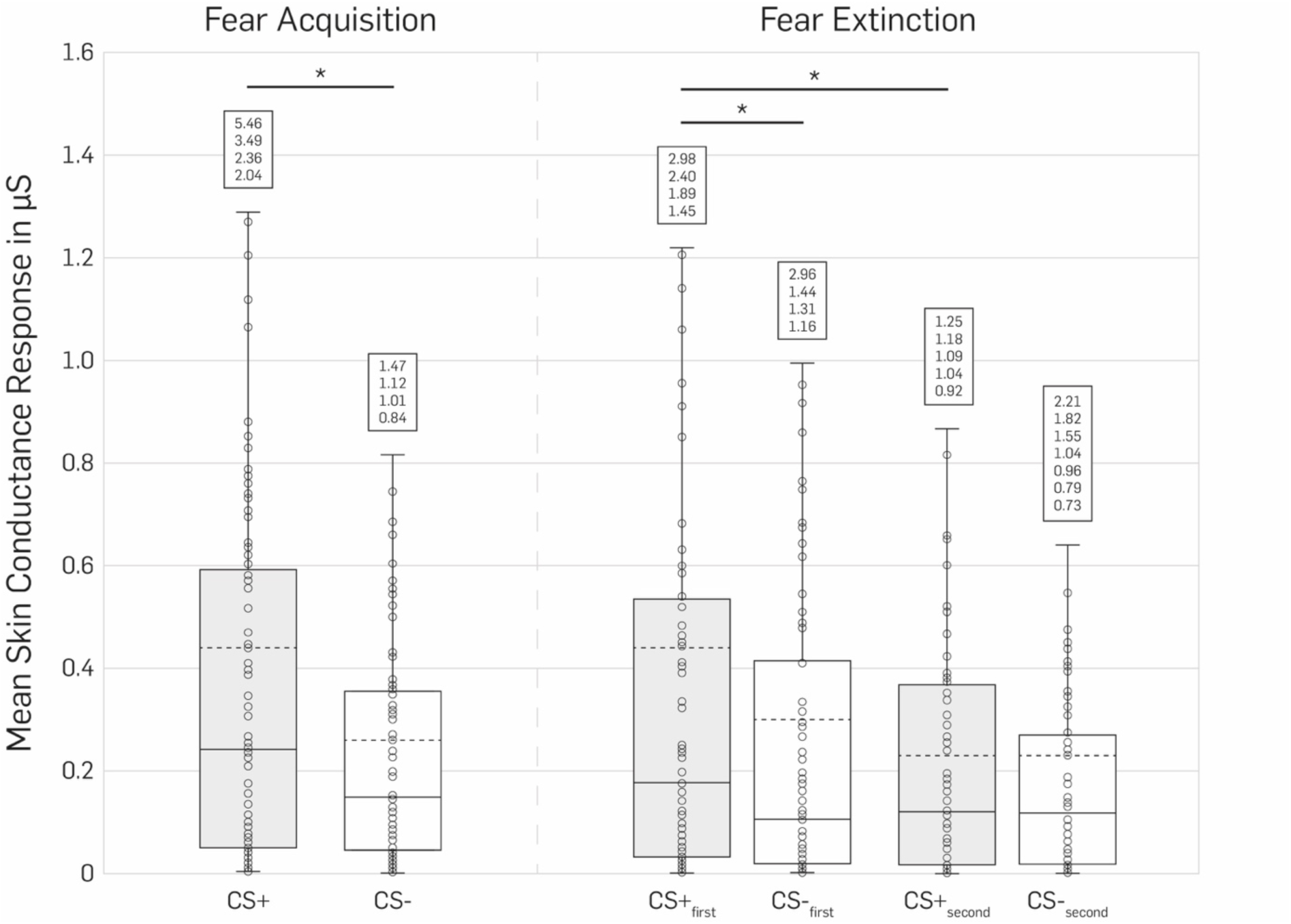
Mean skin conductance responses. Box plots showing the distribution of mean SCRs in reaction to CS+ (grey boxes) and CS-(white boxes) presentations during fear acquisition and extinction. In case of fear acquisition, data were averaged across all 16 CS+ or 16 CS-presentations, respectively. In case of fear extinction, data were averaged across the first or last four CS+ or CS-presentations, respectively. For all plots, mean SCRs are given in microSiemens (µS). For each distribution, the dashed line marks the mean, the solid line marks the median, box edges mark the 25^th^ and 75^th^ percentiles, and whiskers correspond to a 1.5-fold inter-quartile range. Outliers are not shown visually but listed in white boxes for the sake of readability (* p < .01).

### 3.2. SCR-based outcome of fear extinction

SCR values from fear extinction were also averaged across CS+ and CS-trials but separately for the first and second half. These mean values were subjected to a two-way ANOVA with repeated measures. The two factors were “stimulus type” (CS+ vs. CS-) and “time” (first half vs. second half). Results showed a significant interaction effect between “stimulus type” and “time” (F(1,87) = 5.45; *p* < 0.05), indicating that the difference between mean CS+ and CS-values had changed substantially from the first to the second half of fear extinction. As can be seen in Figure 2 (right), mean CS+ values from the first half were higher compared to the corresponding mean CS-values. This difference in mean values almost vanished completely for the second half, likely due to successful fear extinction. As for the fear acquisition data, we also computed one-sample *t*-tests to compare mean CS+ and CS-values from fear extinction. We found that SCR was significantly higher towards CS+ than CS-in the first half (t(87) = 2.651, *p* < .01) but not the second half (t(87) = −0.075, *p* = .941). SCR towards CS+ was significantly higher in the first compared to the second half (t(87) = 2.928, *p* < .01). No significant difference in SCR was observed for CS-between the two halves (t(87) = 1.750, *p* = .084).

### 3.3. Regularized regression models of fear acquisition

Aforementioned measures of fear acquisition and extinction were used as dependent variables in regularized regression models. In each of these regression models, the ROI- or connection-based values which were computed for one out of seven brain variables, served as independent variables. For fear acquisition data, the regularization process yielded six regression models with at least one independent variable exhibiting a relevant association with the SCR-based measure of fear acquisition. There were 135 relevant predictors (out of 70,500 potential predictors) in the RSC model with beta values in the range of -.0873 to .1256 (Supplementary Table 1), 94/70,500 predictors in the DWIC model (βmin = -.0368, βmax = .1243, Supplementary Table 2), 11/360 predictors in the SURF model (βmin = -.0787, βmax = .1003, Supplementary Table 3), 1/360 predictors (R_Pol1, posterior insula) in the THICK model (βmin/max = .0440, Supplementary Table 4), 41/376 predictors in the INVF model (βmin = -.1059, βmax = .1147, Supplementary Table 5), and 2/376 predictors (L_A5, auditory association cortex; L_V3A, visual area V3A) in the ODI model (βmin = -.0633, βmax = -.0302, Supplementary Table 6). In the VOL model, no relevant predictors were left after regularization. The ROIs and connections that exhibited an association in one of these regression models are depicted in Supplementary Figures 2 to 7, either as brain regions on a cortical surface (for predictors from the VOL, THICK, SURF, INVF, and ODI regression models) or as connections in a brain network (for predictors from the RSC and DWIC regression models).

### 3.4. Regularized regression models of fear extinction

The analysis of fear extinction data yielded three regression models that survived the regularization process. There was 1/70,500 predictors (L_putamen - L_LO3, left putamen to lateral occipital area) in the RSC model (β = .0435, Supplementary Table 7), 5/70,500 predictors in the DWIC model (βmin = .0007, βmax = .1274, Supplementary Table 8), and 106/376 predictors in the INVF model (βmin = -.3896, βmax = .2837, Supplementary Table 9). In the VOL, SURF, THICK, and ODI regression models, the regularization process rejected all potential predictors. Relevant predictors of fear extinction are shown in Supplementary Figures 8 to 10.

### 3.5. Predictor overlap among corresponding fear acquisition and extinction models

Some predictors turned out to be relevant in a regression model of fear acquisition as well as the corresponding regression model of fear extinction. The INVF models exhibited a total of eleven such matching ROIs, namely L_2 (primary somatosensory cortex), L_A1 (primary auditory cortex), L_FOP1 (frontal opercular area), L_TPOJ2 (temporo-parieto-occipital junction), L_V3A (visual area V3A), L_V6A (visual area V6A), R_a24pr (part of Brodmann area 24), R_AAIC (anterior agranular insular cortex), R_LBelt (lateral belt of auditory cortex), R_LO2 (lateral occipital area), and R_OP1 (operculum parietale). For the DWIC models, we observed one matching connection, between L_TE1m (part of middle temporal gyrus) and R_p9-46v (part of dorsolateral PFC). In addition, there were four matching network nodes that constituted relevant connections, namely L_TPOJ1 (temporo-parieto-occipital junction), L_p24 (part of Brodmann area 24), R_46 (Brodmann area 46), and R_STV (superior temporal visual area). The RSC models did not show any overlap between fear acquisition and extinction data, neither for whole network connections nor single network nodes.

### 3.6. Predictor overlap among all fear acquisition or extinction models

In our final analysis, we examined the overlap between regression models that had the same dependent variable (SCR-based measure of either fear acquisition or extinction). Specifically, we searched for ROIs (from the VOL, SURF, THICK, INVF, and ODI regression models) and network nodes (from the RSC and DWIC regression models) that were present in two or three different models. In view of regression models based on fear acquisition data, we found 77 ROIs/nodes that replicated across two regression models and 19 ROIs/nodes that replicated across three regression models (Supplementary Tables 10 and 11). Results for the latter were as follows: Eight ROIs/nodes turned out to have a relevant contribution within the RSC, DWIC, and INVF models. These were L_2 (primary somatosensory cortex), L_4 (primary motor cortex), L_8BM (medial part of Brodmann area 8), L_A1 (primary auditory cortex), R_1 (primary somatosensory cortex), R_8BL (lateral part of Brodmann area 8), R_10pp (frontopolar PFC), and R_a24 (anterior cingulate cortex). Five ROIs/nodes repeated across the RSC, DWIC, and SURF models. These were L_PCV (visual posterior region of precuneus), R_FOP1 (frontal opercular area), R_LO1 (lateral occipital area), R_PeEc (perirhinal ectorhinal cortex), and R_PreS (presubiculum). Three ROIs/nodes were present in the RSC, INVF, and SURF models. These were R_MI (middle insular area), R_MIP (medial intraparietal area), and R_PCV (visual posterior region of precuneus). L_A5 (auditory association cortex) was included as a relevant predictor in the RSC, INVF, and ODI models. POI1 (posterior insular area) repeated across the RSC, INVF, and THICK models. R_LBelt (lateral belt of auditory cortex) was present in the DWIC, INVF, and SURF models. These 19 ROIs/nodes are shown in Figure 3 along with the combination of regression models in which they emerged as relevant predictors of fear acquisition.

**Figure 3.**
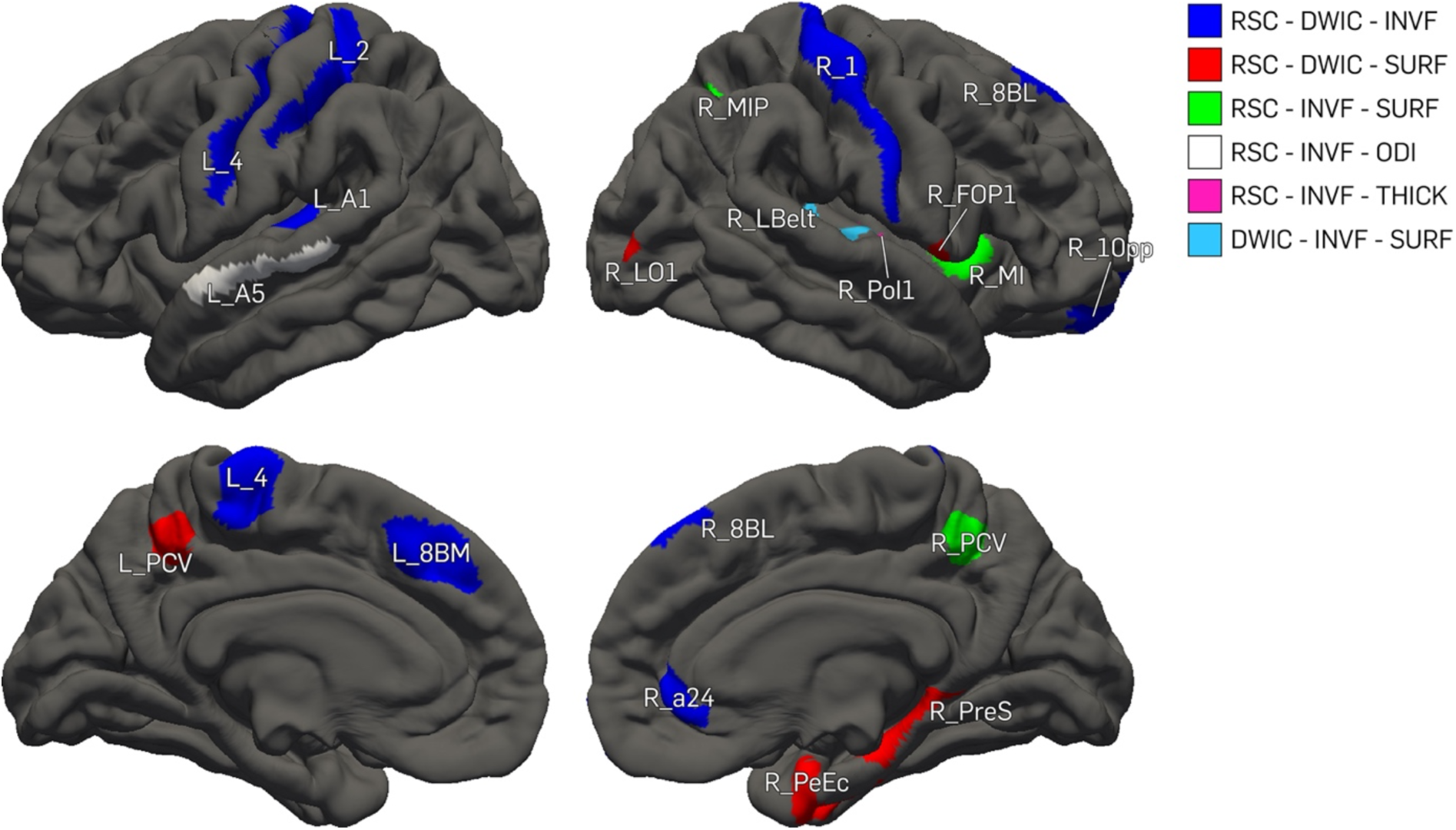
ROIs and network nodes with relevant contributions in three regression models (fear acquisition data). Brain regions are shown on lateral and medial views of cortical surfaces. Each brain region’s color corresponds to a combination of regularized regression models in which respective brain region was included as a relevant predictor, either as a distinct ROI or as a node from a relevant network connection (RSC = resting-state connectivity, DWIC = diffusion-weighted imaging connectivity, INVF = intra-neurite volume fraction, SURF = cortical surface area, ODI = orientation dispersion index, THICK = cortical thickness). Brain regions are labeled according to the Human Connectome Project’s Multimodal Parcellation.

With regard to fear extinction data, we could not identify any predictors that replicated across more than two regression models (Supplementary Table 12). Two regions were identified in the INVF regression model and the DWIC regression model: R_46 (Brodmann area 46) and R_p9-46v (part of dorsolateral PFC) (Figure 4).

**Figure 4.**
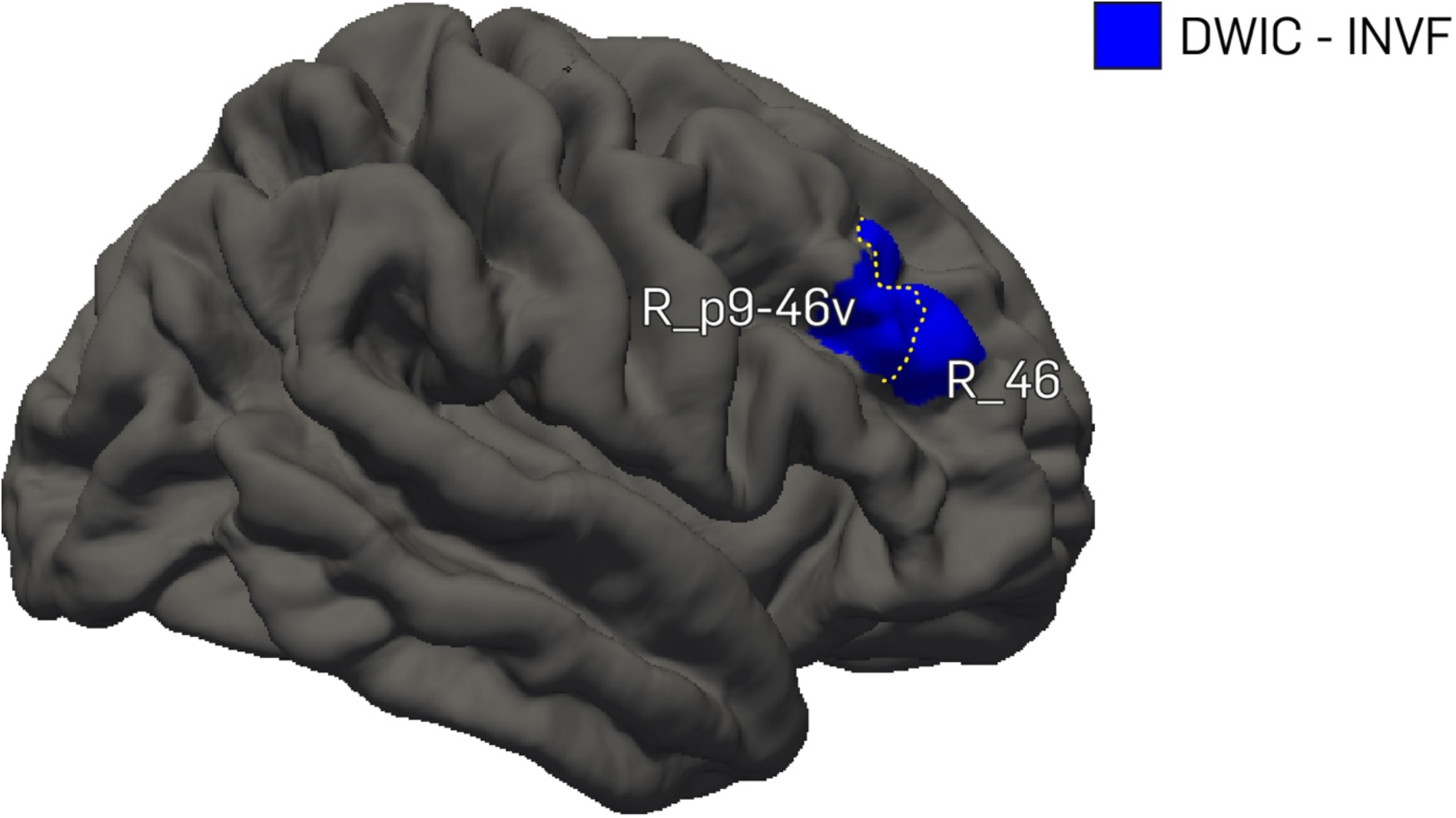
ROIs and network nodes with relevant contributions in two regression models (fear extinction data). Brain regions are shown on a lateral view of the cortical surface. Each brain region’s color corresponds to a combination of regularized regression models in which respective brain region was included as a relevant predictor, either as a distinct ROI or as a node from a relevant network connection (DWIC = diffusion-weighted imaging connectivity, INVF = intra-neurite volume fraction). Brain regions are labeled according to the Human Connectome Project’s Multimodal Parcellation.

## 4 Discussion

In the present study, we related various brain properties to the effect of fear acquisition and extinction, quantified as SCRs towards the CS+ relative to the CS-. These properties included structural connectivity as measured via DWIC, functional connectivity as measured via RSC, brain macrostructure as measured via VOL, THICK, and SURF, as well as brain microstructure as measured via INVF and ODI. We followed a whole-brain approach and did not restrict our analyses to any a-priori specified brain regions. The data-driven nature of this assessment ensured an unbiased analysis of the association of functional and structural properties of individual brain regions and network nodes with fear acquisition and extinction. Using regularized regression models, we found a large number of brain regions and network nodes to be robustly associated with the differentiation between CS+ and CS-during fear acquisition. These were located in the somatosensory, motor, auditory, anterior cingulate, and insular cortices as well as in (para-)hippocampal regions and visual regions of the precuneus. In addition, we observed several regions within the frontal cortex, such as the ventromedial or dorsolateral PFC, that showed significant relationships which replicated across different measures. In contrast, brain regions and network nodes exhibiting multi-modal associations with extinction learning were scarce as only the dorsolateral PFC could be identified as a robust hub.

With respect to the association between fear acquisition and structural/functional brain properties, our results replicate previous findings. For example, various studies have linked the insula and (para-)hippocampus to fear acquisition, both on a functional as well as a structural level (e.g., Fani et al., 2015; Hermans et al., 2017). CS-US contingencies need to be formed and stored in memory for subsequent recall and appropriate behavior whenever the CS+ is presented. In this regard, hippocampal regions are likely to serve as core nodes necessary for the formation of fear memory traces as well as their distribution across the brain. Temporary silencing of or permanent lesions within the hippocampal formation are usually associated with decrements in the recall of acquired fear (for review, see Borgomaneri et al., 2021) indicating its causal role in this process. For the insular cortex, numerous studies have indicated that affective stimuli evoking fear are robustly linked to insula activation (Fullana et al., 2016; Shi et al., 2020). However, emotional processing is mostly associated with activation in the anterior insula whereas our study also revealed robust nodes in the middle and posterior regions of the insula. Especially the posterior insula has been suggested to be fundamentally subserving the processing of painful stimuli (Segerdahl et al., 2015), which could explain the insula’s involvement across its entire anterior-posterior axis in our study. Interestingly, we could not identify any significant nodes in the amygdala. The likelihood of overlapping models was lower for subcortical regions as they were not analyzed across all measures (THICK and SURF were not available). However, we did not find a relationship even at the single model level. At first glance, this finding is surprising given that amygdala activation and connectivity have been linked to the acquisition of fear traces in the past, both by means of correlational and causal methods (e.g., Asok et al., 2018; Knight et al., 2004; LaBar et al., 1998). In contrast to this large body of evidence supporting amygdala involvement in the acquisition of fear, a recent meta-analysis specifically investigating fear conditioning paradigms employing painful stimuli, e.g. electrical stimulation, did not find amygdala activation (Biggs et al., 2020). Hence, it has to be acknowledged that fear conditioning is not inevitably tied to amygdala activation, whereas the activation of insular cortices seems to be more robust. In fact, several meta-as well as mega-analytic findings have cast doubt on the critical involvement of the amygdala in human fear processing (Fullana et al., 2018; Fullana et al., 2016; Visser et al., 2021). Amygdala processing is shown robustly in animal studies that use, for example, foot shocks as their US. In comparison, human fear conditioning paradigms typically do not evoke any behavioral tendencies. Participants will not be inclined to avoid or retreat after experience of shock whereas rodents will immediately try to escape the threatening environment or freeze if escape is not an option (Lamprea et al., 2002). Considering key differences in methodological designs (e.g., Haaker et al., 2019), it is possible that amygdala activation predominantly reflects the processing of threats that are directly linked to behavioral output such as avoidance and flight responses. Other potential reasons as to why there seems to be such a stark discrepancy between human and animal studies with respect to amygdala activation have been proposed to pertain to technical aspects of amygdala delineation, trial-averaging or the lack of intense fear due to sampling healthy cohorts in relatively mildly threatening environments (Fullana et al., 2018).

In accordance with previous meta-analytic findings (Fullana et al., 2016), we could also show robust associations in the anterior cingulate cortex and various parts of the frontal cortex (see Supplementary Table 2). The anterior cingulate cortex is strongly connected to the anterior insular cortex (Cauda et al., 2011) and has been suggested to modulate fear responses (Milad, Quirk, et al., 2007). Direct stimulation or lesioning of the dorsal anterior cingulate cortex has been shown to increase (Mangina & Beuzeron-Mangina, 1996) or decrease (Tranel & Damasio, 1994) autonomic responses as measured via SCRs. Furthermore, the anterior cingulate cortex and the anterior insula are key regions within the salience network (Seeley, 2019) which is involved in the detection of highly salient stimuli such as pain-predicting stimuli presented during a fear conditioning procedure. The involvement of various (pre-)frontal regions such as the ventromedial PFC suggests that successful fear acquisition is accompanied by emotional regulation processes as this region is involved in downregulating activity in fear-associated regions such as the amygdala (Motzkin et al., 2015). Additionally, it has also been proposed that ventromedial PFC activation is likely to reflect the processing of safety signals in contrast to threat signals (Fullana et al., 2016). Assuming that fear learning requires a reliable distinction between threat (CS+) and safety signals (CS-), which is also reflected in the SCR-based difference score we computed to estimate the quality of fear acquisition, a recruitment of regions from the safety network seems highly reasonable.

We also found somatosensory and motor as well as visual and auditory cortices to exhibit robust associations with the effect of fear acquisition. Previous studies have shown that the somatosensory cortex is activated during the perception of painful stimuli (Apkarian et al., 2005). In the present study, the intensity of electrical stimulation was individually set to a level that participants perceived as highly aversive but not painful. However, given that these sensations fall on a continuum and do not represent discrete categories, a relationship between somatosensation and fear acquisition is not surprising. Our results also showed associations between specific properties of the primary motor cortex and the effect of fear acquisition. This is in line with earlier studies (e.g., Butler et al., 2007) reporting an involvement of the primary motor cortex during fear acquisition. Respective activity patterns have been suggested to reflect anticipation of forced movement and tensing of muscles prior to shock delivery via top-down control (Boshuisen et al., 2002). Alternatively, these neural signatures may also represent bottom-up startle responses evoked by the CS+ (Grillon & Baas, 2003). At first glance, the robust associations exhibited by visual regions such as the posterior part of the precuneus could be attributed to the visual nature of the task. However, it has to be noted that detecting the key difference between CS+ and CS-(blue light vs. yellow light) poses a rather basic task, probably not demanding enough to allow for great performance differences. Hence, it seems unlikely that the observed relationships between specific properties of visual brain regions and the effect of fear acquisition can be explained by increased or decreased capability in stimulus distinction. It appears more plausible that respective associations stem from a direct involvement of specific visual regions in the processing of threat-related stimuli. Recently, threat evaluation has been highlighted as a core feature of multiple sensory cortices (for review, Li & Keil, 2023). Here, an initial threat representation in sensory cortices seems to be necessary for fast and precise formation of CS-US associations that are then transmitted to downstream regions for further processing. While associations exhibited by visual brain regions and network nodes can be readily explained in this way, it is more challenging to interpret the involvement of auditory cortex.

Our results revealed a clear discrepancy between the number of brain regions robustly associated with fear acquisition and those involved in fear extinction. We found almost 100 brain regions or network nodes to exhibit multi-modal relationships with fear acquisition and merely two such regions or nodes associated with fear extinction. Previous meta-analytic evidence suggests a considerable overlap between fear acquisition and extinction with respect to functional activation during both conditions. Only the insular, somatosensory, and dorsal anterior cingulate cortices tend to show increased patterns of activity during fear acquisition compared to fear extinction (Fullana et al., 2018). This corroborates our results given that brain regions or network nodes exhibiting robust associations with fear extinction were exclusively observed in the dorsolateral PFC. Fullana et al. (2018) discussed this non-finding in the context of extinction learning being a relatively fragile phenomenon where two memory traces of threat and safety compete against another. Moreover, the authors suggested that the same network could be involved in fear acquisition and extinction while taking over different functions. However, it should be noted that our study used the relative difference between the first and second half of the extinction learning phase and thus directly measured the strength and quality of the extinction learning process in the SCRs. Hence, increased structural connectivity across brain regions, as measured via DWIC, and stronger connections within the dorsolateral PFC, as measured via INVF, were actually associated with more successful fear extinction. The dorsolateral PFC is well-documented in its role within the executive functioning network (Mansouri et al., 2009) and has been reported to impact extinction learning as well (Sotres-Bayon et al., 2006). Extinction learning has been demonstrated to constitute a secondary inhibitory learning that downregulates the initial acquisition memory trace through the involvement of executive functioning (Bouton, 2004). Our findings are clearly in line with this hypothesis.

The current work is subject to several limitations that should be acknowledged. In view of our results concerning fear acquisition, it must be noted that specificity was relatively low as almost half of all potential brain regions were involved in at least one of the employed measures. While this could be due to the large number of measures, eight in total, a more detailed look revealed that most clusters were characterized by a co-occurrence of relevant functional and structural connectivity measures which is unlikely to be incidental. Since our results for the extinction learning phase were highly specific, it also seems unlikely that the analysis approach was overly liberal. As highlighted by Fullana et al. (2018), fear acquisition seems to involve a robust and widespread network across brain regions. Still, future studies are required to replicate the present findings.

In conclusion, we could find a broad range of robust network nodes within the brain to be associated with fear acquisition comprising, among others, the somatosensory, insular, cingulate, and frontal cortices. On the other hand, the effect of extinction learning was exclusively associated with robust activation of the dorsolateral PFC suggesting that neural signatures of extinction learning are less broadly distributed across the brain and more specifically related to executive and inhibitory control. Our study is the first to highlight that neural correlates of specific brain regions show some degree of overlap in their association with fear acquisition and extinction. This finding raises more detailed questions. Future studies may focus on the exact interplay between the neural properties analyzed in our study. They may investigate how the structural and functional connections relevant for fear conditioning are intertwined with each other and how they are affected by the brain’s macro- and microstructural properties (or vice versa). Unraveling the intricate neural correlates of fear learning is imperative not only for expanding our fundamental knowledge of brain function but also for its potential impact on clinical applications. As we continue to delve deeper into the neural circuits and mechanisms that govern fear acquisition and extinction, we are better poised to translate empirical findings into practical interventions. These may include techniques such as individualized brain stimulation used to effectively treat anxiety disorders and related pathologies. In this pursuit, more research is needed to bring about a profound transformation in the lives of those who grapple with the debilitating effects of fear and anxiety.

## Author Contributions

**Christoph Fraenz:** Formal analysis, Investigation, Data Curation, Writing - Original Draft, Writing - Review & Editing, Visualization **Dorothea Metzen:** Formal analysis, Investigation, Data Curation, Writing - Original Draft, Writing - Review & Editing **Julian Packheiser:** Writing - Original Draft, Writing - Review & Editing **Christian J. Merz:** Methodology, Writing - Review & Editing **Helene Selpien:** Investigation, Writing - Review & Editing **Nikolai Axmacher:** Conceptualization, Writing - Review & Editing, Supervision, Project administration **Erhan Genç:** Conceptualization, Methodology, Writing - Review & Editing, Supervision, Project administration

## Data Availability

Data and code used in the analyses are available from the corresponding author upon request.

## Funding and Acknowledgements

This work is part of the Collaborative Research Center SFB 1280 on extinction learning (projects A02, A03, A09, and F02). It was supported by the Deutsche Forschungsgemeinschaft (DFG; project number 31680338). The authors would like to thank PHILIPS Germany (Burkhard Mädler) for scientific support with the MRI measurements as well as Tobias Otto for technical support. The authors would also like to express their gratitude to all research assistants for their support with data acquisition.

## References

Abend, R., Gold, A. L., Britton, J. C., Michalska, K. J., Shechner, T., Sachs, J. F., Winkler, A. M., Leibenluft, E., Averbeck, B. B., & Pine, D. S. (2020). Anticipatory Threat Responding: Associations With Anxiety, Development, and Brain Structure. Biol Psychiatry, 87(10), 916–925. 10.1016/j.biopsych.2019.11.006

Andersson, J. L. R., Graham, M. S., Zsoldos, E., & Sotiropoulos, S. N. (2016). Incorporating outlier detection and replacement into a non-parametric framework for movement and distortion correction of diffusion MR images. Neuroimage, 141, 556–572. 10.1016/j.neuroimage.2016.06.058

Andersson, J. L. R., & Sotiropoulos, S. N. (2016). An integrated approach to correction for off-resonance effects and subject movement in diffusion MR imaging. Neuroimage, 125, 1063–1078. 10.1016/j.neuroimage.2015.10.019

Apkarian, A. V., Bushnell, M. C., Treede, R. D., & Zubieta, J. K. (2005). Human brain mechanisms of pain perception and regulation in health and disease. Eur J Pain, 9(4), 463–484. 10.1016/j.ejpain.2004.11.001

Asok, A., Draper, A., Hoffman, A. F., Schulkin, J., Lupica, C. R., & Rosen, J. B. (2018). Optogenetic silencing of a corticotropin-releasing factor pathway from the central amygdala to the bed nucleus of the stria terminalis disrupts sustained fear. Mol Psychiatry, 23(4), 914–922. 10.1038/mp.2017.79

Bach, D. R., Daunizeau, J., Friston, K. J., & Dolan, R. J. (2010). Dynamic causal modelling of anticipatory skin conductance responses. Biological Psychology, 85(1), 163–170. 10.1016/j.biopsycho.2010.06.007

Behrens, T. E., Berg, H. J., Jbabdi, S., Rushworth, M. F. S., & Woolrich, M. W. (2007). Probabilistic diffusion tractography with multiple fibre orientations: What can we gain? Neuroimage, 34(1), 144–155.

Behrens, T. E., Johansen-Berg, H., Woolrich, M. W., Smith, S. M., Wheeler-Kingshott, C. A., Boulby, P. A., Barker, G. J., Sillery, E. L., Sheehan, K., Ciccarelli, O., Thompson, A. J., Brady, J. M., & Matthews, P. M. (2003). Non-invasive mapping of connections between human thalamus and cortex using diffusion imaging. Nature Neuroscience, 6(7), 750–757.

Biggs, E. E., Timmers, I., Meulders, A., Vlaeyen, J. W. S., Goebel, R., & Kaas, A. L. (2020). The neural correlates of pain-related fear: A meta-analysis comparing fear conditioning studies using painful and non-painful stimuli. Neurosci Biobehav Rev, 119, 52–65. 10.1016/j.neubiorev.2020.09.016

Billiet, T., Vandenbulcke, M., Madler, B., Peeters, R., Dhollander, T., Zhang, H., Deprez, S., van den Bergh, B. R. H., Sunaert, S., & Emsell, L. (2015). Age-related microstructural differences quantified using myelin water imaging and advanced diffusion MRI. Neurobiology of Aging, 36(6), 2107–2121.

Borgomaneri, S., Battaglia, S., Sciamanna, G., Tortora, F., & Laricchiuta, D. (2021). Memories are not written in stone: Re-writing fear memories by means of non-invasive brain stimulation and optogenetic manipulations. Neurosci Biobehav Rev, 127, 334–352. 10.1016/j.neubiorev.2021.04.036

Boshuisen, M. L., Ter Horst, G. J., Paans, A. M., Reinders, A. A., & den Boer, J. A. (2002). rCBF differences between panic disorder patients and control subjects during anticipatory anxiety and rest. Biol Psychiatry, 52(2), 126–135. 10.1016/s0006-3223(02)01355-0

Bouton, M. E. (2004). Context and behavioral processes in extinction. Learn Mem, 11(5), 485–494. 10.1101/lm.78804

Buchel, C., Dolan, R. J., Armony, J. L., & Friston, K. J. (1999). Amygdala-hippocampal involvement in human aversive trace conditioning revealed through event-related functional magnetic resonance imaging. J Neurosci, 19(24), 10869–10876. 10.1523/JNEUROSCI.19-24-10869.1999

Butler, T., Pan, H., Tuescher, O., Engelien, A., Goldstein, M., Epstein, J., Weisholtz, D., Root, J. C., Protopopescu, X., Cunningham-Bussel, A. C., Chang, L., Xie, X. H., Chen, Q., Phelps, E. A., Ledoux, J. E., Stern, E., & Silbersweig, D. A. (2007). Human fear-related motor neurocircuitry. Neuroscience, 150(1), 1–7. 10.1016/j.neuroscience.2007.09.048

Cacciaglia, R., Pohlack, S. T., Flor, H., & Nees, F. (2015). Dissociable roles for hippocampal and amygdalar volume in human fear conditioning. Brain Structure & Function, 220(5), 2575–2586. 10.1007/s00429-014-0807-8

Cauda, F., D’Agata, F., Sacco, K., Duca, S., Geminiani, G., & Vercelli, A. (2011). Functional connectivity of the insula in the resting brain. Neuroimage, 55(1), 8–23. 10.1016/j.neuroimage.2010.11.049

Cohen, M. X., Schoene-Bake, J. C., Elger, C. E., & Weber, B. (2009). Connectivity-based segregation of the human striatum predicts personality characteristics. Nature Neuroscience, 12(1), 32–34.

Craske, M. G., Hermans, D. E., & Vansteenwegen, D. E. (2006). Fear and learning: From basic processes to clinical implications. American Psychological Association.

Daducci, A., Canales-Rodriguez, E. J., Zhang, H., Dyrby, T. B., Alexander, D. C., & Thiran, J. P. (2015). Accelerated microstructure imaging via convex optimization (AMICO) from diffusion MRI data. Neuroimage, 105, 32–44.

Daducci, A., Dal Palu, A., Lemkaddem, A., & Thiran, J. P. (2015). COMMIT: Convex optimization modeling for microstructure informed tractography. IEEE Trans Med Imaging, 34(1), 246–257. 10.1109/TMI.2014.2352414

Dale, A. M., Fischl, B., & Sereno, M. I. (1999). Cortical surface-based analysis. I. Segmentation and surface reconstruction. Neuroimage, 9(2), 179–194.

Fani, N., King, T. Z., Brewster, R., Srivastava, A., Stevens, J. S., Glover, E. M., Norrholm, S. D., Bradley, B., Ressler, K. J., & Jovanovic, T. (2015). Fear-potentiated startle during extinction is associated with white matter microstructure and functional connectivity. Cortex, 64, 249–259. 10.1016/j.cortex.2014.11.006

Feng, P., Feng, T. Y., Chen, Z. C., & Lei, X. (2014). Memory consolidation of fear conditioning: Bi-stable amygdala connectivity with dorsal anterior cingulate and medial prefrontal cortex. Soc Cogn Affect Neurosci, 9(11), 1730–1737. Doi 10.1093/Scan/Nst170

Feng, P., Zheng, Y., & Feng, T. (2016). Resting-state functional connectivity between amygdala and the ventromedial prefrontal cortex following fear reminder predicts fear extinction. Soc Cogn Affect Neurosci, 11(6), 991–1001. 10.1093/scan/nsw031

Feng, T. Y., Feng, P., & Chen, Z. C. (2013). Altered resting-state brain activity at functional MRI during automatic memory consolidation of fear conditioning. Brain Research, 1523, 59–67. Doi 10.1016/J.Brainres.2013.05.039

Fischl, B., Salat, D. H., Busa, E., Albert, M., Dieterich, M., Haselgrove, C., van der Kouwe, A., Killiany, R., Kennedy, D., Klaveness, S., Montillo, A., Makris, N., Rosen, B., & Dale, A. M. (2002). Whole brain segmentation: automated labeling of neuroanatomical structures in the human brain. Neuron, 33(3), 341–355. http://www.ncbi.nlm.nih.gov/pubmed/11832223

http://ac.els-cdn.com/S089662730200569X/1-s2.0-S089662730200569X-main.pdf?_tid=d34b4e74-b11d-11e4-9288-00000aacb361&acdnat=1423570418_d6aa7c50ad5ceb197b47b9781c6f15f7

Fischl, B., Sereno, M. I., & Dale, A. M. (1999). Cortical surface-based analysis. II: Inflation, flattening, and a surface-based coordinate system. Neuroimage, 9(2), 195-207.

Fisher, R. A. (1921). On the probable error of a coefficient of correlation deduced from a small sample. Metron, 1, 3–32.

Fraenz, C., Schlüter, C., Friedrich, P., Jung, R. E., Güntürkün, O., & Genç, E. (2021). Interindividual differences in matrix reasoning are linked to functional connectivity between brain regions nominated by Parieto-Frontal Integration Theory. Intelligence, 87, 101545.

Froeling, M., Tax, C. M., Vos, S. B., Luijten, P. R., & Leemans, A. (2016). ”MASSIVE” brain dataset: Multiple acquisitions for standardization of structural imaging validation and evaluation. Magnetic Resonance in Medicine, 77(5), 1797–1809.

Fullana, M. A., Albajes-Eizagirre, A., Soriano-Mas, C., Vervliet, B., Cardoner, N., Benet, O., Radua, J., & Harrison, B. J. (2018). Fear extinction in the human brain: A meta-analysis of fMRI studies in healthy participants. Neurosci Biobehav Rev, 88, 16–25. 10.1016/j.neubiorev.2018.03.002

Fullana, M. A., Albajes-Eizagirre, A., Soriano-Mas, C., Vervliet, B., Cardoner, N., Benet, O., Radua, J., & Harrison, B. J. (2019). Amygdala where art thou? Neurosci Biobehav Rev, 102, 430–431. 10.1016/j.neubiorev.2018.06.003

Fullana, M. A., Harrison, B. J., Soriano-Mas, C., Vervliet, B., Cardoner, N., Avila-Parcet, A., & Radua, J. (2016). Neural signatures of human fear conditioning: an updated and extended meta-analysis of fMRI studies. Molecular psychiatry, 21(4), 500–508. 10.1038/mp.2015.88

Genc, E., Fraenz, C., Schlueter, C., Friedrich, P., Hossiep, R., Voelkle, M. C., Ling, J. M., Guentuerkuen, O., & Jung, R. E. (2018). Diffusion markers of dendritic density and arborization in gray matter predict differences in intelligence. Nature Communications, 9(1), 1–11.

Genç, E., Metzen, D., Fraenz, C., Schlüter, C., Voelkle, M. C., Arning, L., Streit, F., Nguyen, H. P., Güntürkün, O., Ocklenburg, S., & Kumsta, R. (2023). Structural architecture and brain network efficiency link polygenic scores to intelligence. Human Brain Mapping, n/a(n/a). 10.1002/hbm.26286

Glasser, M. F., Coalson, T. S., Robinson, E. C., Hacker, C. D., Harwell, J., Yacoub, E., Ugurbil, K., Andersson, J., Beckmann, C. F., Jenkinson, M., Smith, S. M., & Van Essen, D. C. (2016). A multi-modal parcellation of human cerebral cortex. Nature, 536(7615), 171–178.

Gongora, D., Vega-Hernandez, M., Jahanshahi, M., Valdes-Sosa, P. A., Bringas-Vega, M. L., & Chbmp. (2020). Crystallized and fluid intelligence are predicted by microstructure of specific white-matter tracts. Hum Brain Mapp, 41(4), 906–916. 10.1002/hbm.24848

Grillon, C., & Baas, J. (2003). A review of the modulation of the startle reflex by affective states and its application in psychiatry. Clin Neurophysiol, 114(9), 1557–1579. 10.1016/s1388-2457(03)00202-5

Haaker, J., Maren, S., Andreatta, M., Merz, C. J., Richter, J., Richter, S. H., Meir Drexler, S., Lange, M. D., Jungling, K., Nees, F., Seidenbecher, T., Fullana, M. A., Wotjak, C. T., & Lonsdorf, T. B. (2019). Making translation work: Harmonizing cross-species methodology in the behavioural neuroscience of Pavlovian fear conditioning. Neurosci Biobehav Rev, 107, 329–345. 10.1016/j.neubiorev.2019.09.020

Hartley, C. A., Fischl, B., & Phelps, E. A. (2011). Brain Structure Correlates of Individual Differences in the Acquisition and Inhibition of Conditioned Fear. Cerebral Cortex, 21(9), 1954–1962. Doi 10.1093/Cercor/Bhq253

Hermann, A., Stark, R., Blecker, C. R., Milad, M. R., & Merz, C. J. (2017). Brain structural connectivity and context-dependent extinction memory. Hippocampus, 27(8), 883–889. 10.1002/hipo.22738

Hermans, E. J., Kanen, J. W., Tambini, A., Fernandez, G., Davachi, L., & Phelps, E. A. (2017). Persistence of Amygdala-Hippocampal Connectivity and Multi-Voxel Correlation Structures During Awake Rest After Fear Learning Predicts Long-Term Expression of Fear. Cereb Cortex, 27(5), 3028–3041. 10.1093/cercor/bhw145

Im, J. J., Kim, B., Hwang, J., Kim, J. E., Kim, J. Y., Rhie, S. J., Namgung, E., Kang, I., Moon, S., Lyoo, I. K., Park, C. H., & Yoon, S. (2017). Diagnostic potential of multimodal neuroimaging in posttraumatic stress disorder. PLoS ONE, 12(5), e0177847. 10.1371/journal.pone.0177847

Jespersen, S. N., Bjarkam, C. R., Nyengaard, J. R., Chakravarty, M. M., Hansen, B., Vosegaard, T., Ostergaard, L., Yablonskiy, D., Nielsen, N. C., & Vestergaard-Poulsen, P. (2010). Neurite density from magnetic resonance diffusion measurements at ultrahigh field: Comparison with light microscopy and electron microscopy. Neuroimage, 49(1), 205–216.

Jespersen, S. N., Leigland, L. A., Cornea, A., & Kroenke, C. D. (2012). Determination of axonal and dendritic orientation distributions within the developing cerebral cortex by diffusion tensor imaging. IEEE Transactions on Medical Imaging, 31(1), 16–32.

Kleckner, I. R., Jones, R. M., Wilder-Smith, O., Wormwood, J. B., Akcakaya, M., Quigley, K. S., Lord, C., & Goodwin, M. S. (2018). Simple, Transparent, and Flexible Automated Quality Assessment Procedures for Ambulatory Electrodermal Activity Data. IEEE Trans Biomed Eng, 65(7), 1460–1467. 10.1109/TBME.2017.2758643

Knight, D. C., Smith, C. N., Cheng, D. T., Stein, E. A., & Helmstetter, F. J. (2004). Amygdala and hippocampal activity during acquisition and extinction of human fear conditioning. Cogn Affect Behav Neurosci, 4(3), 317–325. 10.3758/cabn.4.3.317

Kunimatsu, A., Yasaka, K., Akai, H., Kunimatsu, N., & Abe, O. (2020). MRI findings in posttraumatic stress disorder. J Magn Reson Imaging, 52(2), 380–396. 10.1002/jmri.26929

LaBar, K. S., Gatenby, J. C., Gore, J. C., LeDoux, J. E., & Phelps, E. A. (1998). Human amygdala activation during conditioned fear acquisition and extinction: a mixed-trial fMRI study. Neuron, 20(5), 937–945. 10.1016/s0896-6273(00)80475-4

Lamprea, M. R., Cardenas, F. P., Vianna, D. M., Castilho, V. M., Cruz-Morales, S. E., & Brandao, M. L. (2002). The distribution of fos immunoreactivity in rat brain following freezing and escape responses elicited by electrical stimulation of the inferior colliculus. Brain Res, 950(1-2), 186–194. 10.1016/s0006-8993(02)03036-6

Leemans, A., Jeurissen, B., Sijbers, J., & Jones, D. (2009). ExploreDTI: a graphical toolbox for processing, analyzing, and visualizing diffusion MR data. Proc Intl Soc Mag Reson Med.

Li, W., & Keil, A. (2023). Sensing fear: fast and precise threat evaluation in human sensory cortex. Trends Cogn Sci, 27(4), 341–352. 10.1016/j.tics.2023.01.001

Lonsdorf, T. B., Menz, M. M., Andreatta, M., Fullana, M. A., Golkar, A., Haaker, J., Heitland, I., Hermann, A., Kuhn, M., Kruse, O., Meir Drexler, S., Meulders, A., Nees, F., Pittig, A., Richter, J., Romer, S., Shiban, Y., Schmitz, A., Straube, B.,…Merz, C. J. (2017). Don’t fear ‘fear conditioning’: Methodological considerations for the design and analysis of studies on human fear acquisition, extinction, and return of fear. Neurosci Biobehav Rev, 77, 247–285. 10.1016/j.neubiorev.2017.02.026

Lonsdorf, T. B., & Merz, C. J. (2017). More than just noise: Inter-individual differences in fear acquisition, extinction and return of fear in humans - Biological, experiential, temperamental factors, and methodological pitfalls. Neurosci Biobehav Rev, 80, 703–728. 10.1016/j.neubiorev.2017.07.007

Mangina, C. A., & Beuzeron-Mangina, J. H. (1996). Direct electrical stimulation of specific human brain structures and bilateral electrodermal activity. Int J Psychophysiol, 22(1-2), 1–8. 10.1016/0167-8760(96)00022-0

Mansouri, F. A., Tanaka, K., & Buckley, M. J. (2009). Conflict-induced behavioural adjustment: a clue to the executive functions of the prefrontal cortex. Nature Reviews Neuroscience, 10(2), 141–152. 10.1038/nrn2538

Martynova, O., Tetereva, A., Balaev, V., Portnova, G., Ushakov, V., & Ivanitsky, A. (2020). Longitudinal changes of resting-state functional connectivity of amygdala following fear learning and extinction. Int J Psychophysiol, 149, 15–24. 10.1016/j.ijpsycho.2020.01.002

Milad, M. R., & Quirk, G. J. (2012). Fear Extinction as a Model for Translational Neuroscience: Ten Years of Progress. Annual Review of Psychology, Vol 63, 63, 129–151. 10.1146/annurev.psych.121208.131631

Milad, M. R., Quirk, G. J., Pitman, R. K., Orr, S. P., Fischl, B., & Rauch, S. L. (2007). A role for the human dorsal anterior cingulate cortex in fear expression. Biological Psychiatry, 62(10), 1191–1194. http://ac.els-cdn.com/S0006322307004015/1-s2.0-S0006322307004015-main.pdf?_tid=0f9522ac-ddc3-11e4-ba3c-00000aab0f01&acdnat=1428479237_583233df58899c4c55a39d1f1c09e4b1

Milad, M. R., Wright, C. I., Orr, S. P., Pitman, R. K., Quirk, G. J., & Rauch, S. L. (2007). Recall of fear extinction in humans activates the ventromedial prefrontal cortex and hippocampus in concert. Biological Psychiatry, 62(5), 446–454. Doi 10.1016/J.Biopsych.2006.10.011

Motzkin, J. C., Philippi, C. L., Wolf, R. C., Baskaya, M. K., & Koenigs, M. (2015). Ventromedial prefrontal cortex is critical for the regulation of amygdala activity in humans. Biol Psychiatry, 77(3), 276–284. 10.1016/j.biopsych.2014.02.014

Nees, F., Pohlack, S. T., Grimm, O., Winkelmann, T., Zidda, F., & Flor, H. (2019). White matter correlates of contextual pavlovian fear extinction and the role of anxiety in healthy humans. Cortex, 121, 179–188. 10.1016/j.cortex.2019.08.020

Ocklenburg, S., Friedrich, P., Fraenz, C., Schlueter, C., Beste, C., Gunturkun, O., & Genc, E. (2018). Neurite architecture of the planum temporale predicts neurophysiological processing of auditory speech. Science Advances, 4(7). 10.1126/sciadv.aar6830

Oldfield, R. C. (1971). The assessment and analysis of handedness: The Edinburgh inventory. Neuropsychologia, 9(1), 97–113.

Phelps, E. A. (2004). Human emotion and memory: interactions of the amygdala and hippocampal complex. Current Opinion in Neurobiology, 14(2), 198–202. 10.1016/j.conb.2004.03.015

Pohlack, S. T., Nees, F., Liebscher, C., Cacciaglia, R., Diener, S. J., Ridder, S., Woermann, F. G., & Flor, H. (2012). Hippocampal but not amygdalar volume affects contextual fear conditioning in humans. Hum Brain Mapp, 33(2), 478–488. 10.1002/hbm.21224

Schmitz, J., Fraenz, C., Schluter, C., Friedrich, P., Jung, R. E., Gunturkun, O., Genc, E., & Ocklenburg, S. (2019). Hemispheric asymmetries in cortical gray matter microstructure identified by neurite orientation dispersion and density imaging. Neuroimage, 189, 667–675. 10.1016/j.neuroimage.2019.01.079

Schultz, D. H., Balderston, N. L., & Helmstetter, F. J. (2012). Resting-state connectivity of the amygdala is altered following Pavlovian fear conditioning. Frontiers in Human Neuroscience, 6. Artn 242 Doi 10.3389/Fnhum.2012.00242

Seeley, W. W. (2019). The Salience Network: A Neural System for Perceiving and Responding to Homeostatic Demands. J Neurosci, 39(50), 9878–9882. 10.1523/JNEUROSCI.1138-17.2019

Segerdahl, A. R., Mezue, M., Okell, T. W., Farrar, J. T., & Tracey, I. (2015). The dorsal posterior insula subserves a fundamental role in human pain. Nature Neuroscience, 18(4), 499–500. 10.1038/nn.3969

Sepehrband, F., Alexander, D. C., Kurniawan, N. D., Reutens, D. C., & Yang, Z. (2016). Towards higher sensitivity and stability of axon diameter estimation with diffusion-weighted MRI. NMR Biomed, 29(3), 293–308. https://www.ncbi.nlm.nih.gov/pmc/articles/PMC4949708/pdf/NBM-29-293.pdf

Serang, S., Jacobucci, R., Brimhall, K. C., & Grimm, K. J. (2017). Exploratory Mediation Analysis via Regularization. Struct Equ Modeling, 24(5), 733–744. 10.1080/10705511.2017.1311775

Sherif, T., Rioux, P., Rousseau, M.-E., Kassis, N., Beck, N., Adalat, R., Das, S., Glatard, T., & Evans, A. C. (2014). CBRAIN: a web-based, distributed computing platform for collaborative neuroimaging research. Frontiers in neuroinformatics, 8, 54.

Shi, T., Feng, S., Wei, M., & Zhou, W. (2020). Role of the anterior agranular insular cortex in the modulation of fear and anxiety. Brain Res Bull, 155, 174–183. 10.1016/j.brainresbull.2019.12.003

Smith, S. M., Jenkinson, M., Woolrich, M. W., Beckmann, C. F., Behrens, T. E., Johansen-Berg, H., Bannister, P. R., De Luca, M., Drobnjak, I., Flitney, D. E., Niazy, R. K., Saunders, J., Vickers, J., Zhang, Y., De Stefano, N., Brady, J. M., & Matthews, P. M. (2004). Advances in functional and structural MR image analysis and implementation as FSL. Neuroimage, 23 Suppl 1, S208–219. 10.1016/j.neuroimage.2004.07.051

Sotres-Bayon, F., Cain, C. K., & LeDoux, J. E. (2006). Brain mechanisms of fear extinction: historical perspectives on the contribution of prefrontal cortex. Biol Psychiatry, 60(4), 329–336. 10.1016/j.biopsych.2005.10.012

Tabbert, K., Merz, C. J., Klucken, T., Schweckendiek, J., Vaitl, D., Wolf, O. T., & Stark, R. (2011). Influence of contingency awareness on neural, electrodermal and evaluative responses during fear conditioning. Soc Cogn Affect Neurosci, 6(4), 495–506. 10.1093/scan/nsq070

Tranel, D., & Damasio, H. (1994). Neuroanatomical correlates of electrodermal skin conductance responses. Psychophysiology, 31(5), 427–438. 10.1111/j.1469-8986.1994.tb01046.x

Visser, R. M., Bathelt, J., Scholte, H. S., & Kindt, M. (2021). Robust BOLD Responses to Faces But Not to Conditioned Threat: Challenging the Amygdala’s Reputation in Human Fear and Extinction Learning. J Neurosci, 41(50), 10278–10292. 10.1523/JNEUROSCI.0857-21.2021

Vos, S. B., Tax, C. M., Luijten, P. R., Ourselin, S., Leemans, A., & Froeling, M. (2017). The importance of correcting for signal drift in diffusion MRI. Magn Reson Med, 77(1), 285–299. 10.1002/mrm.26124

Wen, Z., Pace-Schott, E. F., Lazar, S. W., Rosen, J., Ahs, F., Phelps, E. A., LeDoux, J. E., & Milad, M. R. (2024). Distributed neural representations of conditioned threat in the human brain. Nature Communications, 15(1), 2231. 10.1038/s41467-024-46508-0

Wen, Z., Raio, C. M., Pace-Schott, E. F., Lazar, S. W., LeDoux, J. E., Phelps, E. A., & Milad, M. R. (2022). Temporally and anatomically specific contributions of the human amygdala to threat and safety learning. Proc Natl Acad Sci U S A, 119(26), e2204066119. 10.1073/pnas.2204066119

Winkelmann, T., Grimm, O., Pohlack, S. T., Nees, F., Cacciaglia, R., Dinu-Biringer, R., Steiger, F., Wicking, M., Ruttorf, M., Schad, L. R., & Flor, H. (2016). Brain morphology correlates of interindividual differences in conditioned fear acquisition and extinction learning. Brain Structure & Function, 221(4), 1927–1937. 10.1007/s00429-015-1013-z

Zhang, H., Schneider, T., Wheeler-Kingshott, C. A., & Alexander, D. C. (2012). NODDI: Practical in vivo neurite orientation dispersion and density imaging of the human brain. Neuroimage, 61(4), 1000–1016.

Zou, H., & Hastie, T. (2005). Regularization and variable selection via the elastic net. Journal of the Royal Statistical Society Series B: Statistical Methodology, 67(2), 301–320.

