## Supplementary Table 1 for "Multi-modal brain properties are associated with interindividual differences in fear acquisition and extinction"

**Supplementary Table 1. Beta coefficients from the fear acquisition and RSC regression model.**

| **Connection** | **Beta** |
| --- | --- |
| R_s32 - R_Pir | 0.1256 |
| R_s32 - L_PoI1 | 0.1062 |
| L_V1 - L_PI | 0.0946 |
| R_TPOJ1 - R_PSL | 0.0864 |
| R_Ig - R_EC | 0.0844 |
| L_accumbens-area - R_d23ab | 0.0743 |
| R_AIP - L_VMV2 | 0.0649 |
| R_OP4 - L_PEF | 0.0631 |
| R_Ig - L_OP2-3 | 0.0484 |
| R_STV - R_55b | 0.0466 |
| R_STGa - L_MIP | 0.0464 |
| L_pallidum - R_STSvp | 0.0422 |
| L_MT - L_A5 | 0.0374 |
| R_V3 - L_Pir | 0.0329 |
| L_V4t - L_7PL | 0.0327 |
| L_VMV1 - L_PoI1 | 0.0304 |
| R_s32 - R_PoI1 | 0.0302 |
| R_MBelt - L_IFSp | 0.0292 |
| R_TPOJ1 - R_FOP1 | 0.0291 |
| R_PF - R_FEF | 0.0286 |
| L_PHT - L_11l | 0.0275 |
| R_Ig - R_FOP4 | 0.0269 |
| R_55b - R_45 | 0.0261 |
| R_STSda - L_MT | 0.0253 |
| R_LIPd - R_52 | 0.0205 |
| R_PHA3 - L_OP2-3 | 0.0199 |
| L_VVC - L_Ig | 0.0189 |
| R_8C - R_8BL | 0.0182 |
| L_PeEc - L_A1 | 0.0174 |
| R_7AL - L_A1 | 0.0165 |
| R_TPOJ2 - R_LIPd | 0.0165 |
| R_TPOJ1 - R_FOP4 | 0.0155 |
| R_V4t - R_A1 | 0.0151 |
| R_accumbens-area - R_d23ab | 0.0134 |
| R_47s - L_7PL | 0.0129 |
| R_FOP1 - L_FOP3 | 0.0126 |
| R_IP0 - R_11l | 0.0123 |
| R_PSL - R_55b | 0.0120 |
| R_47s - L_9-46d | 0.0119 |
| R_ventraldc - L_a10p | 0.0107 |
| L_PFt - L_OFC | 0.0102 |
| L_7PL - L_5m | 0.0097 |
| L_STSvp - L_Pir | 0.0094 |
| R_PCV - R_31pv | 0.0055 |
| L_DVT - L_33pr | 0.0051 |
| L_PeEc - L_IFJa | 0.0050 |
| R_IFJa - R_a47r | 0.0049 |
| R_6v - L_OP2-3 | 0.0037 |
| L_POS2 - L_PCV | 0.0036 |
| L_STSdp - L_PGi | 0.0035 |
| R_25 - L_43 | 0.0032 |
| R_6mp - L_VMV1 | 0.0030 |
| R_accumbens-area - L_STSva | 0.0025 |
| L_PHA3 - L_OP2-3 | 0.0023 |
| R_PGp - L_pOFC | 0.0020 |
| R_5mv - L_PHA3 | 0.0013 |
| R_PCV - L_33pr | 0.0012 |
| L_accumbens-area - L_52 | 0.0005 |
| R_TPOJ1 - R_FOP3 | 0.0001 |
| L_IP0 - L_6r | 0.0000 |
| L_TPOJ3 - L_TA2 | 0.0000 |
| R_9a - L_FEF | 0.0000 |
| R_TE1a - R_EC | 0.0000 |
| R_IFJp - L_d32 | 0.0000 |
| R_PeEc - L_9-46d | -0.0002 |
| L_43 - L_23d | -0.0004 |
| R_45 - R_7PL | -0.0005 |
| R_47m - L_10pp | -0.0006 |
| L_V6A - L_PHA2 | -0.0025 |
| R_PGi - L_p9-46v | -0.0025 |
| R_PGi - L_7Pm | -0.0029 |
| R_PGi - L_LIPd | -0.0029 |
| R_PeEc - L_SCEF | -0.0030 |
| R_PEF - R_IPS1 | -0.0040 |
| R_STV - R_i6-8 | -0.0048 |
| R_i6-8 - R_9-46d | -0.0050 |
| R_ventraldc - R_PeEc | -0.0050 |
| R_i6-8 - R_8BM | -0.0051 |
| L_13l - L_6d | -0.0051 |
| R_5m - L_7m | -0.0057 |
| R_TPOJ3 - L_p9-46v | -0.0065 |
| R_ventraldc - R_PGp | -0.0068 |
| R_AAIC - L_TA2 | -0.0075 |
| R_TE1p - R_8BM | -0.0076 |
| R_PGi - L_LIPv | -0.0081 |
| R_IFSa - L_pOFC | -0.0084 |
| R_IPS1 - L_6v | -0.0090 |
| R_a24 - R_10pp | -0.0092 |
| R_10v - L_TGv | -0.0103 |
| R_8Av - L_PEF | -0.0110 |
| L_TPOJ2 - L_10pp | -0.0123 |
| R_11l - L_SFL | -0.0126 |
| R_IFJp - R_d32 | -0.0131 |
| R_TPOJ3 - R_MIP | -0.0140 |
| R_5mv - L_PBelt | -0.0148 |
| R_V6A - L_PIT | -0.0153 |
| R_TE1m - R_47s | -0.0155 |
| R_IPS1 - L_PFcm | -0.0156 |
| R_TGd - R_9a | -0.0156 |
| L_PCV - L_PBelt | -0.0161 |
| R_TE2a - L_IP0 | -0.0176 |
| L_i6-8 - L_8BM | -0.0178 |
| R_44 - L_STSvp | -0.0189 |
| R_PeEc - L_7PC | -0.0193 |
| R_SFL - R_i6-8 | -0.0197 |
| R_Ig - R_6d | -0.0205 |
| L_pOFC - L_6d | -0.0222 |
| L_PCV - L_43 | -0.0227 |
| R_v23ab - R_44 | -0.0233 |
| R_pallidum - R_PHA2 | -0.0263 |
| R_PeEc - L_STSva | -0.0275 |
| L_TE2a - L_OFC | -0.0275 |
| L_p32pr - L_OP4 | -0.0292 |
| R_31a - L_7Pm | -0.0299 |
| L_PFcm - L_a32pr | -0.0300 |
| R_RSC - R_8BM | -0.0310 |
| R_9a - L_10v | -0.0322 |
| L_MI - L_23d | -0.0348 |
| L_31a - L_7Pm | -0.0395 |
| R_PreS - L_p24 | -0.0402 |
| R_7PL - L_5mv | -0.0426 |
| L_PI - L_IFJa | -0.0430 |
| R_FFC - R_A5 | -0.0447 |
| R_LO1 - R_IFJa | -0.0509 |
| R_9a - L_TE2a | -0.0515 |
| L_s32 - L_6d | -0.0520 |
| R_i6-8 - R_31a | -0.0536 |
| L_PCV - L_52 | -0.0549 |
| R_V6A - R_PIT | -0.0575 |
| R_PGi - L_PeEc | -0.0582 |
| R_45 - L_IP0 | -0.0587 |
| L_9-46d - L_6r | -0.0645 |
| R_PGi - L_IP0 | -0.0820 |
| R_9m - L_IP0 | -0.0863 |
| R_Ig - R_10pp | -0.0873 |
