## Supplementary Table 2 for "Multi-modal brain properties are associated with interindividual differences in fear acquisition and extinction"

**Supplementary Table 2. Beta coefficients from the fear acquisition and DWIC regression model.**

| **Connection** | **Beta** |
| --- | --- |
| R_IFSa - L_p24 | 0.1243 |
| R_45 - L_p24 | 0.0866 |
| R_PSL - R_LO1 | 0.0860 |
| R_p24pr - L_44 | 0.0762 |
| L_MIP - L_8Av | 0.0760 |
| R_PH - R_PBelt | 0.0730 |
| R_PFt - R_FOP1 | 0.0725 |
| R_TE2a - R_LO1 | 0.0711 |
| R_STV - R_LO1 | 0.0711 |
| R_PH - R_LO1 | 0.0688 |
| R_Ig - R_FFC | 0.0643 |
| R_V1 - L_MBelt | 0.0495 |
| L_PHA3 - L_8Av | 0.0429 |
| R_LO1 - R_FFC | 0.0417 |
| L_a24pr - L_46 | 0.0405 |
| R_a9-46v - R_10pp | 0.0395 |
| R_p24pr - R_6ma | 0.0372 |
| R_10d - L_9m | 0.0371 |
| L_accumbens-area - R_p10p | 0.0370 |
| R_p24pr - R_p10p | 0.0368 |
| R_a24 - R_46 | 0.0329 |
| R_p24 - R_52 | 0.0324 |
| R_LBelt - L_ProS | 0.0299 |
| R_MT - R_Ig | 0.0298 |
| R_p9-46v - L_TE1m | 0.0287 |
| R_FEF - L_IP1 | 0.0205 |
| R_SCEF - L_s6-8 | 0.0185 |
| R_47l - R_13l | 0.0159 |
| R_RSC - R_7AL | 0.0159 |
| R_TE2p - R_PBelt | 0.0159 |
| R_PBelt - L_ProS | 0.0128 |
| R_V8 - R_a24pr | 0.0125 |
| L_PGs - L_A1 | 0.0119 |
| R_TE2p - R_MBelt | 0.0109 |
| R_V4 - R_A1 | 0.0104 |
| R_FOP1 - R_44 | 0.0096 |
| R_p10p - L_p24 | 0.0094 |
| R_SCEF - L_Ig | 0.0083 |
| L_MIP - L_i6-8 | 0.0066 |
| R_TE2a - R_STSva | 0.0066 |
| L_s6-8 - L_IFSp | 0.0062 |
| R_a9-46v - R_9m | 0.0059 |
| R_9p - L_FOP2 | 0.0058 |
| L_TA2 - L_8BM | 0.0056 |
| R_VMV2 - R_V8 | 0.0055 |
| R_VVC - R_6d | 0.0052 |
| R_p24pr - L_a9-46v | 0.0049 |
| R_PoI2 - R_FOP4 | 0.0047 |
| R_IFJp - L_24dv | 0.0045 |
| L_AVI - L_44 | 0.0043 |
| L_s6-8 - L_PGi | 0.0043 |
| R_IFSa - R_8BL | 0.0040 |
| R_6ma - L_Ig | 0.0027 |
| L_PHA1 - L_24dv | 0.0019 |
| R_a32pr - R_52 | 0.0005 |
| R_TE2p - R_A4 | 0.0004 |
| R_accumbens-area - R_4 | 0.0004 |
| R_6a - L_p32 | 0.0003 |
| R_FOP1 - R_2 | 0.0001 |
| R_LO1 - L_10pp | 0.0000 |
| R_IFSp - R_4 | 0.0000 |
| R_VMV1 - L_13l | -0.0007 |
| R_V6 - R_PeEc | -0.0007 |
| R_PreS - L_AVI | -0.0013 |
| R_V6 - R_PreS | -0.0013 |
| R_V3 - L_9-46d | -0.0013 |
| R_V6 - R_H | -0.0020 |
| R_STSva - L_45 | -0.0021 |
| R_POS2 - R_p32pr | -0.0030 |
| R_V6 - R_DVT | -0.0031 |
| R_V2 - L_6d | -0.0031 |
| R_7Pm - L_AAIC | -0.0036 |
| R_TA2 - L_9-46d | -0.0041 |
| L_PCV - L_MST | -0.0046 |
| R_V4t - L_AAIC | -0.0053 |
| R_EC - L_STSva | -0.0060 |
| L_TPOJ1 - L_FST | -0.0069 |
| R_TA2 - R_3a | -0.0070 |
| R_s32 - L_PBelt | -0.0109 |
| R_V3A - L_9-46d | -0.0112 |
| R_POS2 - R_31pd | -0.0138 |
| L_TF - L_AAIC | -0.0163 |
| R_PreS - R_H | -0.0164 |
| R_accumbens-area - L_8C | -0.0168 |
| R_3b - R_3a | -0.0171 |
| R_LO1 - L_MI | -0.0177 |
| R_6d - L_POS2 | -0.0197 |
| R_accumbens-area - L_8BL | -0.0212 |
| L_caudate - L_9a | -0.0278 |
| R_IPS1 - L_47s | -0.0310 |
| R_a32pr - R_4 | -0.0311 |
| L_IFSp - L_45 | -0.0362 |
| R_TE1a - R_10r | -0.0363 |
| R_p32 - L_10v | -0.0368 |
