## Supplementary Table 3 for "Multi-modal brain properties are associated with interindividual differences in fear acquisition and extinction"

**Supplementary Table 3. Beta coefficients from the fear acquisition and cortical surface area regression model.**

| **ROI** | **Beta** |
| --- | --- |
| R_MIP | 0.1003 |
| R_LBelt | 0.0918 |
| R_FOP1 | 0.0291 |
| L_PCV | 0.0274 |
| R_v23ab | 0.0252 |
| R_LO1 | 0.0144 |
| R_PCV | 0.0021 |
| R_PGp | 0.0018 |
| R_PeEc | -0.0170 |
| R_H | -0.0451 |
| R_PreS | -0.0787 |
