## Supplementary Table 5 for "Multi-modal brain properties are associated with interindividual differences in fear acquisition and extinction"

**Supplementary Table 5. Beta coefficients from the fear acquisition and INVF regression model.**

| **ROI** | **Beta** |
| --- | --- |
| R_8BL | 0.1147 |
| L_V3A | 0.0934 |
| L_8BM | 0.0924 |
| L_TPOJ2 | 0.0837 |
| R_8Av | 0.0835 |
| R_a24pr | 0.0779 |
| R_6ma | 0.0770 |
| R_OP1 | 0.0717 |
| R_5mv | 0.0579 |
| R_AAIC | 0.0405 |
| L_2 | 0.0389 |
| L_3a | 0.0367 |
| R_PoI1 | 0.0277 |
| R_9p | 0.0186 |
| L_9m | 0.0158 |
| R_8Ad | 0.0109 |
| R_LBelt | 0.0108 |
| L_TGv | 0.0106 |
| L_V6A | 0.0080 |
| L_VIP | -0.0029 |
| L_LBelt | -0.0059 |
| R_LO2 | -0.0062 |
| R_MIP | -0.0105 |
| L_46 | -0.0116 |
| R_10v | -0.0177 |
| R_VVC | -0.0205 |
| L_9a | -0.0207 |
| L_7PL | -0.0231 |
| L_A1 | -0.0257 |
| R_10pp | -0.0330 |
| R_pallidum | -0.0335 |
| R_31pv | -0.0376 |
| L_V6 | -0.0499 |
| L_V3B | -0.0596 |
| L_H | -0.0735 |
| R_VMV2 | -0.0749 |
| R_STSdp | -0.0760 |
| L_FOP1 | -0.0811 |
| L_8Ad | -0.0915 |
| L_A5 | -0.0922 |
| R_PCV | -0.1059 |
