## Supplementary Table 8 for "Multi-modal brain properties are associated with interindividual differences in fear acquisition and extinction"

**Supplementary Table 8. Beta coefficients from the fear extinction and DWIC regression model.**

| **Connection** | **Beta** |
| --- | --- |
| R_46 - L_TE1m | 0.1274 |
| R_p9-46v - L_TE1m | 0.0535 |
| R_STV - L_V7 | 0.0410 |
| R_p9-46v - L_TPOJ1 | 0.0055 |
| R_46 - L_p24 | 0.0007 |
