## Supplementary Table 9 for "Multi-modal brain properties are associated with interindividual differences in fear acquisition and extinction"

**Supplementary Table 9. Beta coefficients from the fear extinction and INVF regression model.**

| **ROI** | **Beta** |
| --- | --- |
| R_TE2p | 0.2837 |
| L_TPOJ2 | 0.2712 |
| R_OP1 | 0.2124 |
| R_TGv | 0.1939 |
| R_PHA1 | 0.1762 |
| L_V3A | 0.1715 |
| R_52 | 0.1678 |
| L_4 | 0.1669 |
| L_6v | 0.1584 |
| R_s6-8 | 0.1303 |
| R_TE1p | 0.1302 |
| L_TGd | 0.1299 |
| R_EC | 0.1281 |
| L_47l | 0.1264 |
| R_a24pr | 0.1180 |
| R_7AL | 0.1178 |
| L_47m | 0.1158 |
| R_13l | 0.1139 |
| L_V3 | 0.1029 |
| L_POS1 | 0.1003 |
| L_V6A | 0.0852 |
| R_46 | 0.0745 |
| L_33pr | 0.0653 |
| R_FOP5 | 0.0610 |
| L_TPOJ3 | 0.0568 |
| L_PHA2 | 0.0546 |
| R_AAIC | 0.0535 |
| R_ProS | 0.0525 |
| R_10d | 0.0477 |
| R_OP2-3 | 0.0469 |
| L_OP2-3 | 0.0465 |
| L_PBelt | 0.0459 |
| L_47s | 0.0414 |
| L_PEF | 0.0402 |
| R_IFSa | 0.0362 |
| L_IP2 | 0.0342 |
| R_LO2 | 0.0323 |
| L_IP0 | 0.0311 |
| L_31pv | 0.0304 |
| R_PHT | 0.0290 |
| R_IFJa | 0.0259 |
| R_6r | 0.0197 |
| R_d23ab | 0.0178 |
| L_TE2p | 0.0140 |
| L_24dv | 0.0138 |
| R_TGd | 0.0137 |
| L_V8 | 0.0111 |
| R_Pir | 0.0083 |
| R_PSL | 0.0053 |
| L_TE2a | 0.0044 |
| R_hippocampus | 0.0035 |
| R_STSvp | 0.0024 |
| L_MST | 0.0009 |
| L_Ig | 0.0007 |
| L_PFm | 0.0000 |
| R_LIPd | 0.0000 |
| L_IPS1 | -0.0002 |
| L_IFSp | -0.0002 |
| L_LO1 | -0.0026 |
| R_7PL | -0.0038 |
| L_2 | -0.0043 |
| R_MBelt | -0.0052 |
| L_MIP | -0.0056 |
| R_A5 | -0.0067 |
| L_STSdp | -0.0108 |
| L_pOFC | -0.0124 |
| R_V6A | -0.0133 |
| R_MT | -0.0160 |
| R_V8 | -0.0173 |
| L_FOP3 | -0.0208 |
| R_a32pr | -0.0233 |
| R_TA2 | -0.0261 |
| R_LBelt | -0.0274 |
| R_TPOJ3 | -0.0318 |
| R_RSC | -0.0349 |
| R_31pd | -0.0465 |
| L_25 | -0.0466 |
| R_45 | -0.0475 |
| R_8BM | -0.0482 |
| L_MBelt | -0.0571 |
| L_MI | -0.0582 |
| L_3b | -0.0618 |
| L_A1 | -0.0669 |
| R_9m | -0.0676 |
| R_7m | -0.0683 |
| L_7AL | -0.0686 |
| L_PH | -0.0778 |
| R_PI | -0.0784 |
| R_SCEF | -0.0803 |
| R_p9-46v | -0.0831 |
| R_7Pm | -0.0992 |
| R_47l | -0.0998 |
| R_SFL | -0.1267 |
| L_52 | -0.1268 |
| L_PI | -0.1292 |
| L_caudate | -0.1331 |
| R_p10p | -0.1339 |
| R_1 | -0.1360 |
| R_FST | -0.1416 |
| L_d32 | -0.1480 |
| R_V3A | -0.1604 |
| R_IP0 | -0.1699 |
| L_FOP1 | -0.1740 |
| L_PFop | -0.1988 |
| L_PHA1 | -0.2608 |
| L_PoI2 | -0.3896 |
