## Supplementary Table 10 for "Multi-modal brain properties are associated with interindividual differences in fear acquisition and extinction"

**Supplementary Table 10. ROIs and network nodes with relevant contributions in two regression models (fear acquisition data).**

| **ROI** | **RSC** | **DWIC** | **INVF** | **ODI** | **VOL** | **SURF** | **THICK** |
| --- | --- | --- | --- | --- | --- | --- | --- |
| L_1 | X | X |  |  |  |  |  |
| L_6d | X | X |  |  |  |  |  |
| L_7PL | X |  | X |  |  |  |  |
| L_9-46d | X | X |  |  |  |  |  |
| L_9a |  | X | X |  |  |  |  |
| L_9m |  | X | X |  |  |  |  |
| L_10pp | X | X |  |  |  |  |  |
| L_10v | X | X |  |  |  |  |  |
| L_13l | X | X |  |  |  |  |  |
| L_46 |  | X | X |  |  |  |  |
| L_i6-8 | X | X |  |  |  |  |  |
| L_IFSp | X | X |  |  |  |  |  |
| L_Ig | X | X |  |  |  |  |  |
| L_MI | X | X |  |  |  |  |  |
| L_MIP | X | X |  |  |  |  |  |
| L_p24 | X | X |  |  |  |  |  |
| L_p32 | X | X |  |  |  |  |  |
| L_PBelt | X | X |  |  |  |  |  |
| L_PGi | X | X |  |  |  |  |  |
| L_PH | X | X |  |  |  |  |  |
| L_PHA3 | X | X |  |  |  |  |  |
| L_POS2 | X | X |  |  |  |  |  |
| L_STSva | X | X |  |  |  |  |  |
| L_TA2 | X | X |  |  |  |  |  |
| L_TGv | X |  | X |  |  |  |  |
| L_TPOJ2 | X |  | X |  |  |  |  |
| L_V3 |  |  | X | X |  |  |  |
| L_V3A |  |  | X | X |  |  |  |
| L_V6 | X |  | X |  |  |  |  |
| L_V6A | X |  | X |  |  |  |  |
| R_2 | X | X |  |  |  |  |  |
| R_4 | X | X |  |  |  |  |  |
| R_5m | X |  | X |  |  |  |  |
| R_5mv | X |  | X |  |  |  |  |
| R_6d | X | X |  |  |  |  |  |
| R_6ma |  | X | X |  |  |  |  |
| R_7AL | X | X |  |  |  |  |  |
| R_8Av | X |  | X |  |  |  |  |
| R_9m | X | X |  |  |  |  |  |
| R_9p |  | X | X |  |  |  |  |
| R_10v | X |  | X |  |  |  |  |
| R_31pv | X |  | X |  |  |  |  |
| R_44 | X | X |  |  |  |  |  |
| R_45 | X | X |  |  |  |  |  |
| R_52 | X | X |  |  |  |  |  |
| R_A1 | X | X |  |  |  |  |  |
| R_a24pr |  | X | X |  |  |  |  |
| R_AAIC | X |  | X |  |  |  |  |
| R_EC | X | X |  |  |  |  |  |
| R_FEF | X | X |  |  |  |  |  |
| R_FFC | X | X |  |  |  |  |  |
| R_FOP4 | X | X |  |  |  |  |  |
| R_H |  | X |  |  |  | X |  |
| R_IFJp | X | X |  |  |  |  |  |
| R_IFSa | X | X |  |  |  |  |  |
| R_Ig | X | X |  |  |  |  |  |
| R_IPS1 | X | X |  |  |  |  |  |
| R_MBelt | X | X |  |  |  |  |  |
| R_PF | X | X |  |  |  |  |  |
| R_PGp | X |  |  |  |  | X |  |
| R_PH | X | X |  |  |  |  |  |
| R_PSL | X | X |  |  |  |  |  |
| R_RSC | X | X |  |  |  |  |  |
| R_s32 | X | X |  |  |  |  |  |
| R_STV | X | X |  |  |  |  |  |
| R_TE1a | X | X |  |  |  |  |  |
| R_TE2a | X | X |  |  |  |  |  |
| R_V3 | X | X |  |  |  |  |  |
| R_V4 | X | X |  |  |  |  |  |
| R_V4t | X | X |  |  |  |  |  |
| R_V6 | X | X |  |  |  |  |  |
| R_v23ab | X |  |  |  |  | X |  |
| R_VMV2 |  | X | X |  |  |  |  |
| R_VVC |  | X | X |  |  |  |  |
| lh_accumbens-area | X | X |  |  |  |  |  |
| rh_accumbens-area | X | X |  |  |  |  |  |
| rh_pallidum | X |  | X |  |  |  |  |

RSC = resting-state connectivity, DWIC = diffusion weighted imaging connectivity, INVF = intra-neurite volume fraction, ODI = orientation dispersion index, VOL = regional brain volume, SURF = cortical surface area, THICK = cortical thickness
