## Supplementary Table 11 for "Multi-modal brain properties are associated with interindividual differences in fear acquisition and extinction"

**Supplementary Table 11. ROIs and network nodes with relevant contributions in three regression models (fear acquisition data).**

| **ROI** | **RSC** | **DWIC** | **INVF** | **ODI** | **VOL** | **SURF** | **THICK** |
| --- | --- | --- | --- | --- | --- | --- | --- |
| L_2 | X | X | X |  |  |  |  |
| L_4 | X | X | X |  |  |  |  |
| L_8BM | X | X | X |  |  |  |  |
| L_A1 | X | X | X |  |  |  |  |
| L_A5 | X |  | X | X |  |  |  |
| L_PCV | X | X |  |  |  | X |  |
| R_1 | X | X | X |  |  |  |  |
| R_8BL | X | X | X |  |  |  |  |
| R_10pp | X | X | X |  |  |  |  |
| R_a24 | X | X | X |  |  |  |  |
| R_FOP1 | X | X |  |  |  | X |  |
| R_LBelt |  | X | X |  |  | X |  |
| R_LO1 | X | X |  |  |  | X |  |
| R_MI | X |  | X |  |  | X |  |
| R_MIP | X |  | X |  |  | X |  |
| R_PCV | X |  | X |  |  | X |  |
| R_PeEc | X | X |  |  |  | X |  |
| R_PoI1 | X |  | X |  |  |  | X |
| R_PreS | X | X |  |  |  | X |  |

RSC = resting-state connectivity, DWIC = diffusion weighted imaging connectivity, INVF = intra-neurite volume fraction, ODI = orientation dispersion index, VOL = regional brain volume, SURF = cortical surface area, THICK = cortical thickness
