## Supplementary Table 12 for "Multi-modal brain properties are associated with interindividual differences in fear acquisition and extinction"

**Supplementary Table 12. ROIs and network nodes with relevant contributions in two regression models (fear extinction data).**

| **ROI** | **RSC** | **DWIC** | **INVF** | **ODI** | **VOL** | **SURF** | **THICK** |
| --- | --- | --- | --- | --- | --- | --- | --- |
| R_46 |  | X | X |  |  |  |  |
| R_p9-46v |  | X | X |  |  |  |  |

RSC = resting-state connectivity, DWIC = diffusion weighted imaging connectivity, INVF = intra-neurite volume fraction, ODI = orientation dispersion index, VOL = regional brain volume, SURF = cortical surface area, THICK = cortical thickness
